# Replicate libraries set the noise floor for cell-state claims in cross-sectional single-cell data: an empirical audit in drug-tolerant EGFR-mutant PC9 cells

**DOI:** 10.64898/2026.11.19.767931

**Authors:** Wasim Raza, Saba Mehmood, Carsten Enderlein, Fabrício Alves Barbosa Da Silva

**Affiliations:** Developmental Biology and Dynamical Systems Group, Universidade Federal do Rio de Janeiro, Campus Duque de Caxias Professor Geraldo Cidade, Duque de Caxias, Rio de Janeiro, Brazil; Department of Mathematics, University of Management and Technology (UMT), Lahore, Pakistan; NUMPEX-Nano, Campus Duque de Caxias da UFRJ, Rio de Janeiro, Brazil; Scientific Computing Programme, Oswaldo Cruz Foundation (FIOCRUZ), Rio de Janeiro, Brazil; Department of Mathematics, Faculty of Sciences, Thal University, Bhakkar, Pakistan

**Keywords:** single-cell RNA-seq, replicate libraries, batch effects, drug-tolerant persister cells, cell-state attractors, energy distance, lineage barcodes, PC9, EGFR

## Abstract

**Background:** Claims that drug-tolerant cancer cells occupy stable states rest on single-cell snapshots with one sequencing library per condition, where a difference between conditions cannot be separated from a difference between libraries. No standard says how far a contrast must exceed replicate-library variation.

**Results:** We contribute a replicate-referenced framework: a distance ratio expressing a contrast as a multiple of the distance between replicate libraries of one condition, and a library-level permutation test giving exact ***p*** values by relabelling whole libraries. In simulations, single-library tests and a split-half reference rejected a true null in 99 to 100% of runs, and Harmony followed by a cell-level test did so in 12 to 32% with structured library effects. With four libraries per arm, the library-level test held false positives at 3 to 5% in every setting. In public osimertinib-treated PC9 data with two libraries per condition (GSE150949), halves of one library were indistinguishable, replicate libraries separated at 59 to 93% accuracy, and contrasts between times in drug and sort gates exceeded replicate distance 4.5- to 51-fold. In Drop-seq erlotinib data with one library per condition (GSE149383), two libraries of the same condition separated more readily (93.3%) than withdrawn and re-treated cells (88.5%).

**Conclusions:** Cross-sectional contrasts should be reported as multiples of replicate-library distance and, with four or more libraries per arm, tested by library-level permutation. Cells moved away from the untreated profile and stayed away, but the data do not support attractor-style inference, nor do they show that tolerant states are not bistable.

## Background

Cancer cells that survive targeted therapy are often described as occupying stable, non-genetic states, and the evidence is mostly single-cell snapshots in which each condition is one sequencing library. In that design a difference between conditions cannot be told apart from a difference between libraries, and how far a contrast exceeds the difference between independent libraries of one condition decides how much of it can be read as biology. The question matters in epidermal growth factor receptor (*EGFR*) mutant non-small-cell lung cancer, where tyrosine kinase inhibitors (TKIs) such as osimertinib, a standard first-line choice [1], produce marked regressions that are rarely durable. A minority of tumour cells survives drug doses that kill the bulk population, and this fraction can seed overt resistance [2, 3]. These drug-tolerant persisters were reported to be reversible after drug withdrawal and to depend on a non-mutational chromatin configuration [4], and the phenomenon has since been described for other tumour types and drugs [5–7]. Re-treatment with the same EGFR-TKI after an intervening therapy gives only modest responses (pooled objective response rate 15% and disease control rate 61% across nineteen heterogeneous studies [8]), which raises the question of whether tolerance is fully reversible.

A dynamical reading of this state-based view treats tolerant phenotypes as stable states, or attractors, of a gene-regulatory network. It follows Waddington’s landscape [9], Kauffman’s Boolean networks [10] and the formalisation of stable phenotypes as attractors [11–13]. Experimental work includes sorted breast cancer subpopulations that relax to a fixed equilibrium mixture [14], rare pre-existing resistance-competent melanoma cells [15], lineage barcoding towards resistance [16], cycling persisters that arise from restricted pre-existing lineages [17] and resistant phenotypes that arise from non-genetic transitions [18] (reviewed in [19, 20]). Support for attractors specifically remains largely qualitative [21], and separated clusters on a uniform manifold approximation and projection (UMAP) plot are a common form of evidence.

Library-to-library variation is a recognised problem in single-cell work [22–24], and several tools address neighbouring questions. Batch-mixing metrics such as kBET quantify how well libraries mix [25]. Pseudobulk and mixed-model approaches treat the sample and not the cell as the unit of replication [26, 27]. Classifier separability ranks cell types by their response to a perturbation [28], and neighbourhood-based tests of differential abundance use replicates [29]. Batch-correction methods such as Harmony remove library structure before comparison [30], and distance-based multivariate tests with a nested design compare variation between groups with variation within them [31, 32]. These approaches assume that replicate libraries exist and are used to test or remove differences between conditions. What persister studies lack is an empirical scale on which a contrast is expressed as a multiple of the difference between replicate libraries of one condition, together with evidence on which comparisons control false positives when library effects are present.

### Closest prior work and what is new here

Three bodies of work come closest, and none answers the question we ask. First, replicate designs have been used to measure technical variation: Tung and colleagues processed replicate collections of induced pluripotent stem cell lines and showed that batch effects are substantial and can be estimated only with replicates [33], but they studied untreated lines and gave no scale for judging a contrast between conditions. Second, benchmarks of multi-sample differential-state methods showed that treating cells as independent replicates inflates false positives and that sample-level methods perform better [26, 27, 34, 35]. These benchmarks ask which genes differ when replicates exist, and they do not include single-library or split-half references, whole-distribution contrasts or an exact test across libraries. Third, theory shows that snapshots alone cannot fix cell dynamics without further assumptions [36], but this is an argument in principle and not a measurement in a drug-tolerance system. The persister studies whose data we reuse [17, 37] did not set out to quantify replicate variation as a reference for state contrasts. To our knowledge, no earlier study has (i) measured replicate-library variation in a drug-tolerant cancer cell system and expressed state contrasts as multiples of it, (ii) benchmarked single-library, split-half, held-out-library, batch-corrected, pseudobulk and library-level permutation approaches against one another under library effects of known structure, or (iii) shown in a public persister dataset that two libraries of one condition separate as readily as a drug-history contrast. Permuting whole libraries rests on a classical exchangeability argument, and we do not present that principle as new. The original contributions are the replicate-referenced distance ratio, the calibrated library-level test applied to Harmony-corrected cells, the benchmark that shows which comparisons fail, and the empirical audit of two public PC9 datasets.

We address this in three steps. First, we define a replicate-referenced distance ratio, *ρ_ED_*, that expresses the energy distance between two conditions [32] as a multiple of the distance between replicate libraries of one condition. We pair it with three controls: split halves of one library, classification of a held-out replicate, and comparisons inside a library using lineage barcodes. We also define a library-level permutation test (LPT), which calibrates any statistic that does not use arm labels exactly, by relabelling whole libraries. Second, we benchmark these in simulations of known truth against single-library classifiers and distance tests, a split-half reference, held-out-library classification, Harmony followed by a cell-level test, pseudobulk testing (our own moderated-*t* implementation and DESeq2) and Augur. The simulations include low-rank library effects, changes in the occupancy of a tolerant state, and replicate-to-replicate variation in that occupancy, with ablations of the reference, the distance, the number of principal components, cells per library and libraries per arm. Third, we apply the framework to two public datasets of PC9 cells (an *EGFR* exon-19-deletion line). GSE150949 is a 10x osimertinib time course with two replicate libraries per condition [17]. GSE149383 is a Drop-seq erlotinib time course with a withdrawal and re-treatment arm and one library per condition [37].

In simulation, single-library and split-half comparisons rejected the null in nearly every run. Among replicate-aware approaches, only the library-level permutation test kept false positives low in every setting, and applying it to Harmony-corrected cells gave the highest power of the exact tests in most settings. In GSE150949, expression moved away from the untreated profile by far more than the difference between replicate libraries, and cells sharing a lineage barcode were closer than random cells in every library. In GSE149383, two libraries of the same condition separated more readily than the withdrawn and re-treated arms. The Results give the evidence, and the Discussion sets out where the framework applies.

## Results

### Replicate libraries set a noise floor, and halves of one library never separate

GSE150949 contains 11 libraries from one experiment. These are Day 0 (one library), Day 3 and Day 7 (two replicate libraries each), and Day 14, sorted on a proliferation reporter into high-, medium- and low-retention gates with two replicate libraries per gate [17]. All 56,419 cells passed quality control, depth was uncorrelated with PC1 after regression (*r* = 0.000), and we used the full 2,000-gene panel (Methods). We compared libraries at three levels, always at the same sample size (1,300 cells per group, 20 PCs and 20 random draws). The results are in Fig 1a,b and Table 1. Randomly splitting one library into halves gave accuracies of 0.497 to 0.505 and energy distances of −0.003 to 0.002 in all 11 libraries, so the classifier and the distance showed no detectable bias. Two replicate libraries of the same condition were nevertheless separable, with 93.0% accuracy at Day 3, 88.0% at Day 7, 85.3% for the low-retention Day 14 gate and 64.4% and 59.0% for the high- and medium-retention gates (energy distance 0.10 to 1.67). A replicate effect is therefore present even in a carefully designed 10x experiment, and it varies widely between conditions. Replicate cultures differ in biology as well as technical handling, so we call it a replicate effect and do not attribute it to either. The Day 3 value (93%) is similar to the 93.3% we find between two Day 11 libraries of the Drop-seq dataset below. The platforms, protocols and sample sizes (195 cells per library there) differ, so we draw no inference from the resemblance.

**Fig. 1.**
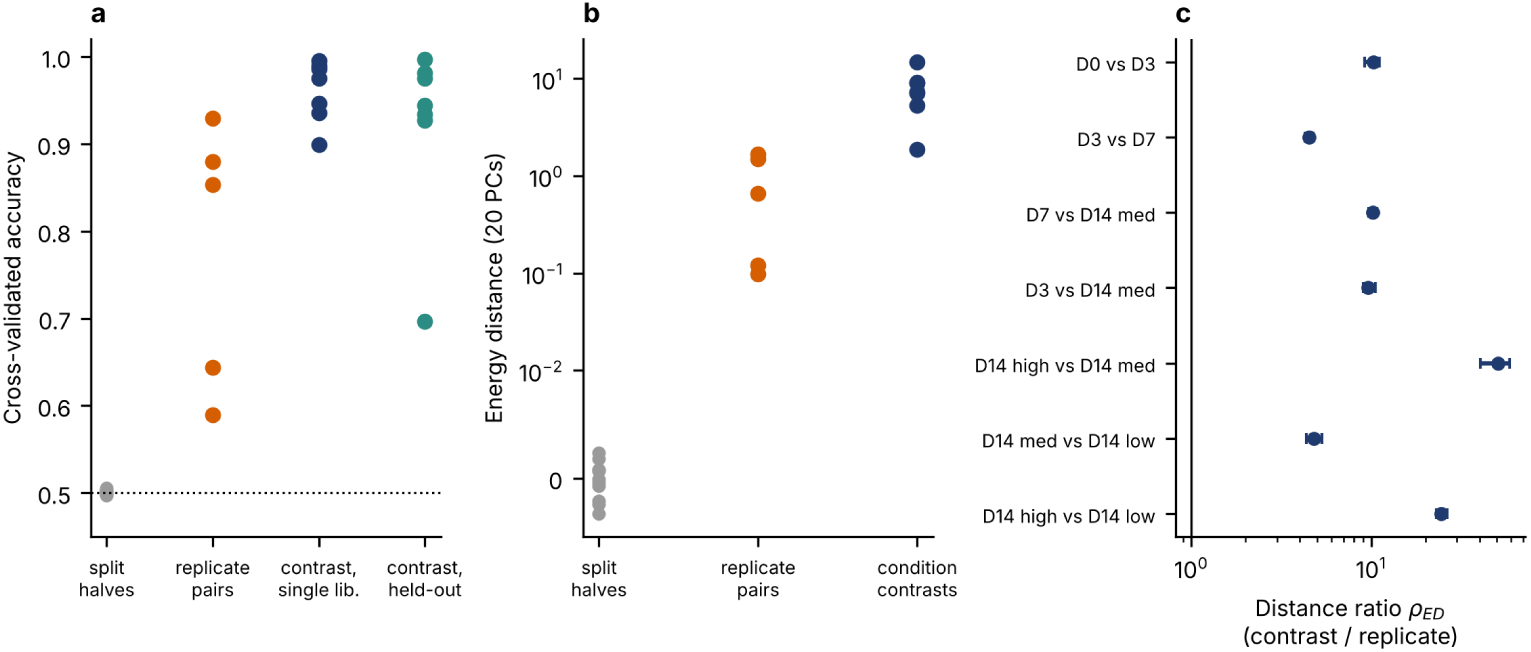
Condition contrasts against the replicate noise floor in GSE150949. **a**, Cross-validated accuracy (1,300 cells per group, 20 PCs) for halves of one library (grey, 11 libraries), replicate libraries of one condition (orange, five conditions), condition contrasts with one library per arm (blue) and condition contrasts classified with a held-out replicate (green, seven contrasts). Dotted line, chance. **b**, Energy distance on a symmetric logarithmic axis for the same three levels. **c**, Distance ratio *ρ_ED_* for the seven contrasts (mean and 2.5 to 97.5% range of 20 subsampling draws). Vertical line, ratio of 1.

**Table 1.** Noise floor in GSE150949 from halves of one library and from replicate libraries.

| Comparison | Cells (library 1 / 2) | Accuracy | Energy distance |
|---|---|---|---|
| Split halves of one library (all 11 libraries) | n/a | 0.497 to 0.505 | −0.003 to 0.002 |
| Replicate libraries, Day 3 | 7,093 / 7,273 | 0.930 | 1.50 |
| Replicate libraries, Day 7 | 4,770 / 3,515 | 0.880 | 1.67 |
| Replicate libraries, Day 14, high retention | 5,380 / 5,449 | 0.644 | 0.10 |
| Replicate libraries, Day 14, medium retention | 4,221 / 5,097 | 0.590 | 0.12 |
| Replicate libraries, Day 14, low retention | 2,690 / 4,153 | 0.853 | 0.66 |

1,300 cells per group (matched), 20 PCs, mean of 20 random draws. Accuracy is from five-fold cross-validated logistic regression, and chance is 0.5. Energy distance has expectation 0 for identical distributions.

The replicate difference was neither confined to one direction nor spread evenly. The direction of the centroid difference alone separated replicate libraries with 0.56 to 0.82 accuracy, which is 58 to 82% of the above-chance accuracy of the 20-PC classifier 4 (0.59 to 0.93). Between 18 and 52% of the squared centroid shift lay in PC1 to PC3 (Additional file 3).

### Condition contrasts clear the floor, and the distance ratio shows by how much

We compared seven contrasts (Table 2). With one library per arm every contrast was separable (90 to 99.6%), which is what a single-library design would report. The replicate-referenced accuracy ratio *ρ* = (acc_contrast_ − 0.5)*/*(acc_replicate_ − 0.5) was 1.2 to 3.9. Accuracy is saturated by the replicate effect itself (replicate accuracy 0.61 to 0.93) and so understates the gap. The energy distance between conditions was 4.5 to 51 times the distance between replicate libraries (*ρ_ED_*, Fig 1c). The smallest ratios were for Day 3 versus Day 7 (4.5, range 4.3 to 4.8) and for medium versus low retention at Day 14 (4.8, range 4.4 to 5.3). Classifiers trained on one replicate and tested on the other classified 92.7 to 99.7% of cells correctly for six of the seven contrasts and 69.7% for medium versus low retention, the contrast that includes the most variable replicate pair. Time in drug and sort gate therefore produce differences that exceed replicate variation. For contrasts with Day 0, which is a single library, the comparison is with the Day 3 replicate distance. Explaining the Day 0 versus Day 3 distance by a Day 0 library effect would require that effect to be about ten times the difference seen between the two Day 3 libraries.

**Table 2.**
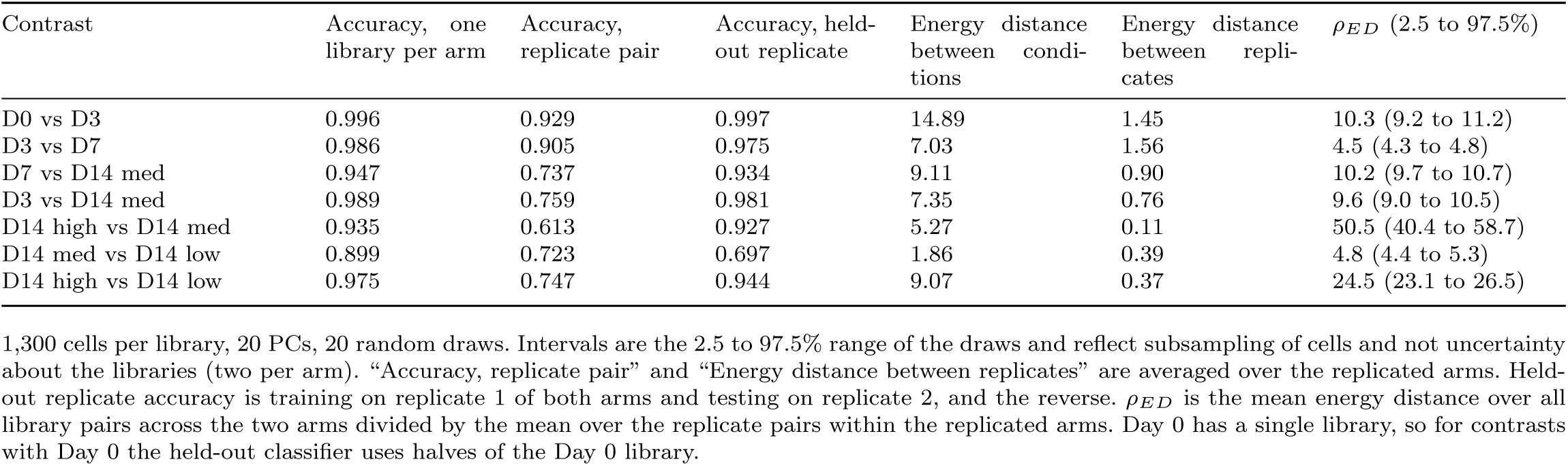
Condition contrasts in GSE150949 against the replicate noise floor.

| Contrast | Accuracy, one library per arm | Accuracy, replicate pair | Accuracy, held-out replicate | Energy distance between condi-tions | Energy distance between repli-cates | ρ<sub>ED</sub> (2.5 to 97.5%) |
|---|---|---|---|---|---|---|
| D0 vs D3 | 0.996 | 0.929 | 0.997 | 14.89 | 1.45 | 10.3 (9.2 to 11.2) |
| D3 vs D7 | 0.986 | 0.905 | 0.975 | 7.03 | 1.56 | 4.5 (4.3 to 4.8) |
| D7 vs D14 med | 0.947 | 0.737 | 0.934 | 9.11 | 0.90 | 10.2 (9.7 to 10.7) |
| D3 vs D14 med | 0.989 | 0.759 | 0.981 | 7.35 | 0.76 | 9.6 (9.0 to 10.5) |
| D14 high vs D14 med | 0.935 | 0.613 | 0.927 | 5.27 | 0.11 | 50.5 (40.4 to 58.7) |
| D14 med vs D14 low | 0.899 | 0.723 | 0.697 | 1.86 | 0.39 | 4.8 (4.4 to 5.3) |
| D14 high vs D14 low | 0.975 | 0.747 | 0.944 | 9.07 | 0.37 | 24.5 (23.1 to 26.5) |

1,300 cells per library, 20 PCs, 20 random draws. Intervals are the 2.5 to 97.5% range of the draws and reflect subsampling of cells and not uncertainty about the libraries (two per arm). “Accuracy, replicate pair” and “Energy distance between replicates” are averaged over the replicated arms. Held-out replicate accuracy is training on replicate 1 of both arms and testing on replicate 2, and the reverse. ρ<sub>ED</sub> is the mean energy distance over all library pairs across the two arms divided by the mean over the replicate pairs within the replicated arms. Day 0 has a single library, so for contrasts with Day 0 the held-out classifier uses halves of the Day 0 library.

### Replicate pairs are nearest neighbours, and the fold change depends on the number of PCs

The seven contrasts above were chosen by biological interest. To check that they do not drive the picture, we computed the energy distance between all 11 libraries (Fig 2a; 800 cells per library, 5 draws, 20 PCs, computed on the processed principal components of GSE150949). For each of the 10 libraries that has a replicate, the nearest of the other 10 libraries was its own replicate, at 10, 20 and 50 PCs alike. Day 0, which has no replicate, was closest to a medium-retention Day 14 library. Replicate distances at 20 PCs were 1.4 (Day 3), 1.7 (Day 7), 0.1 (high retention), 0.1 (medium) and 0.7 (low). Distances between libraries of different conditions ranged from 1.2 to 16.3. The weakest margin was medium versus low retention at Day 14, where the closest cross-condition library pair (1.2) was only 1.8 times the larger of the two replicate distances (0.66). This agrees with its lower held-out accuracy (69.7%, Table 2).

**Fig. 2.**
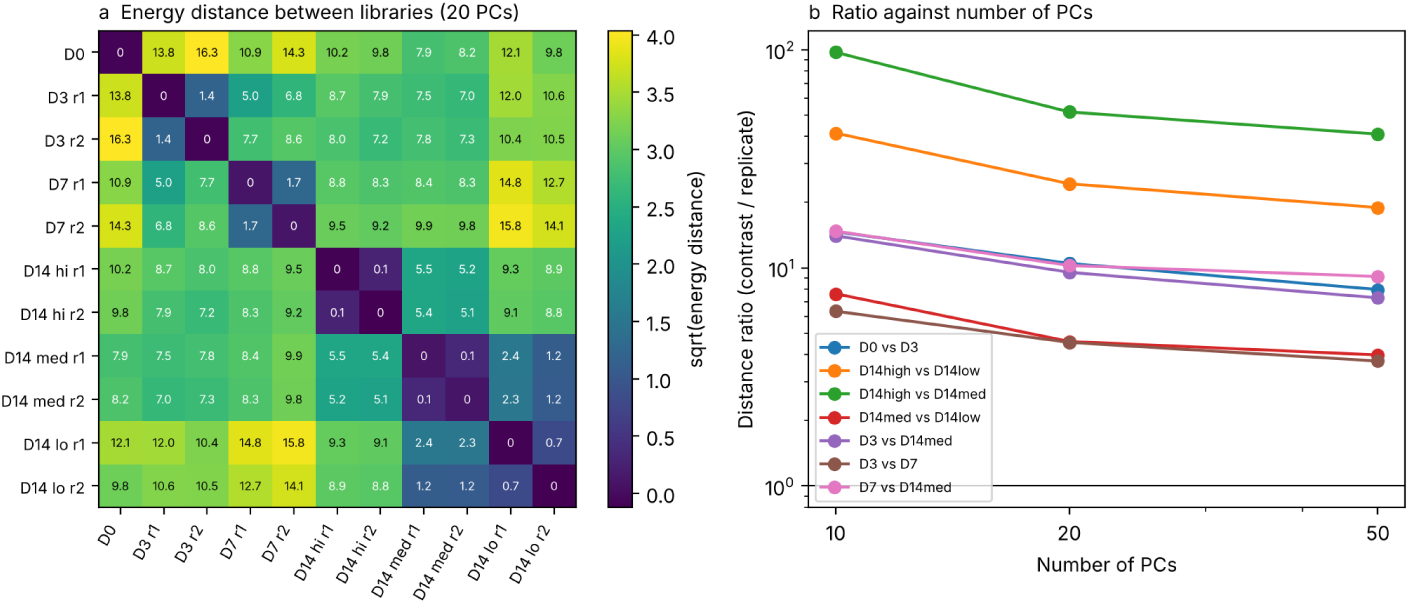
All-library distance matrix and sensitivity of the distance ratio to the number of PCs in GSE150949. **a**, Energy distance between all 11 libraries (20 PCs, 800 cells per library, mean of 5 draws). The diagonal is the split-half distance of each library. Colour shows the square root of the distance and numbers show the distance. r1 and r2 are replicate libraries. **b**, Distance ratio for the seven contrasts at 10, 20 and 50 PCs (logarithmic axes). Horizontal line, ratio of 1.

The distance ratio fell as more PCs were used (Table 3 and Fig 2b), because replicate differences are relatively larger in the higher-dimensional space. All seven ratios stayed at or above 3.7 from 10 to 50 PCs, so the qualitative conclusion holds. The fold values should be read as orders of magnitude and not as estimates with known precision. The largest ratios (high versus medium retention, 41 to 97) come from a denominator of about 0.1 that is estimated from one library pair per arm, and the figure of 51 reported above is therefore the least stable number in this paper.

**Table 3.** Sensitivity of the energy-distance ratio in GSE150949 to the number of PCs.

| Contrast | 10 PCs | 20 PCs | 50 PCs | Closest library pair / largest replicate distance (20 PCs) |
|---|---|---|---|---|
| D0 vs D3 | 14.6 | 10.5 | 8.0 | 9.6 |
| D3 vs D7 | 6.3 | 4.5 | 3.7 | 3.0 |
| D7 vs D14 med | 14.7 | 10.2 | 9.1 | 5.0 |
| D3 vs D14 med | 14.0 | 9.5 | 7.3 | 4.9 |
| D14 high vs D14 med | 97.3 | 51.8 | 41.0 | 44.0 |
| D14 med vs D14 low | 7.6 | 4.6 | 4.0 | 1.8 |
| D14 high vs D14 low | 41.5 | 24.3 | 18.9 | 13.4 |

Ratio of the mean energy distance over all library pairs across the two arms to the mean over the replicate pairs within the replicated arms. 800 cells per library, mean of 5 draws, fixed seed. Values at 20 PCs differ slightly from Table 2 (1,300 cells, 20 draws). For contrasts with Day 0 the denominator is the Day 3 replicate distance. The last column compares the single closest pair of libraries across the two arms with the larger replicate distance. Day 0 is one library.

A second caution comes from the simulation (Additional file 2). The replicate difference in these data is concentrated and not diffuse. The direction of the centroid difference alone recovers 58 to 82% of the above-chance accuracy of the 20-PC classifier, and up to 52% of the squared centroid shift lies in PC1 to PC3. In simulations with low-rank library effects and two libraries per arm, the null distribution of the ratio was heavy-tailed (95th percentile 3.2 to 3.6). That threshold belongs to the simulation and is not transferable to these data. It does mean that the two smallest ratios here (Day 3 versus Day 7 and medium versus low retention, 4.5 and 4.8 at 20 PCs, 3.7 and 4.0 at 50) clear a heavy-tailed null only narrowly, and that a library-level test, which needs four or more libraries per arm and is not possible here, would be needed before they are read as strong evidence. The larger ratios are not threatened by this.

### A one-dimensional summary of trajectory shape cannot be resolved against replicate variation

The mean distance of cells from the Day 0 centroid rose from 11.3 at Day 0 to 20.2 and 22.0 in the two Day 3 libraries (Fig 3a and Table 4). This is a rise of 9.8 units on average, about five times the largest replicate difference (2.0). The change between Day 3 and Day 7 (21.1 to 19.7, a fall of 1.4) and between Day 7 and the Day 14 gates (19.7 to 20.4, a rise of 0.7) was smaller than the replicate differences at Day 3 (1.8), Day 7 (2.0) and the low-retention Day 14 gate (1.9). A rise, fall and rise shape drawn from one library per time point would in single-library data be read as overshoot and partial relaxation. Along this axis it cannot be resolved. This statement concerns one summary of the data and not the populations themselves. The Day 3 and Day 7 libraries are clearly different populations (distance ratio 4.5, held-out accuracy 97.5%, Table 2), so expression did change between those time points. What these data cannot establish is the size and direction of that change along the distance-from-Day-0 axis.

**Fig. 3.**
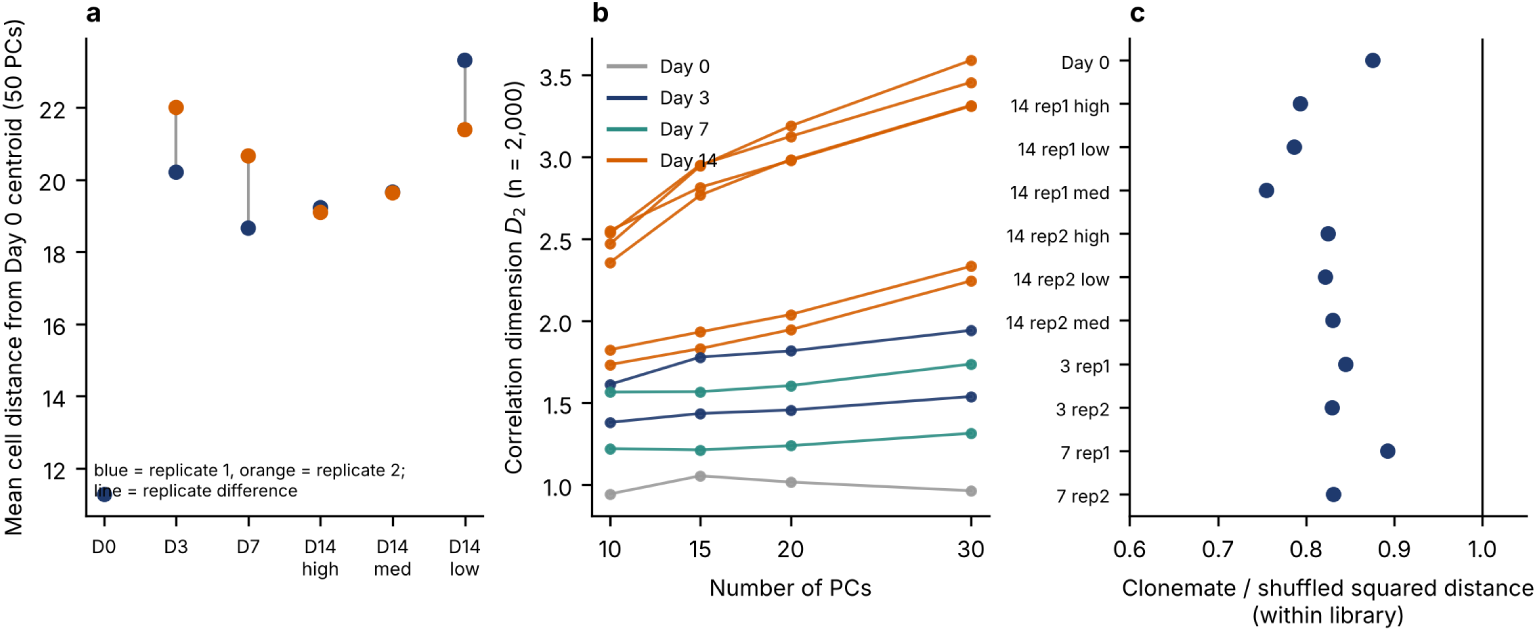
Trajectory, correlation dimension and lineage concordance in GSE150949. **a**, Mean cell distance from the Day 0 centroid (50 PCs) for each library, and the two replicates of a condition are joined by a line. **b**, Correlation dimension *D*_2_ (2,000 cells, mean of 8 draws) against the number of PCs for each library, coloured by time point. **c**, Within-library squared distance between cells sharing a lineage barcode divided by the mean squared distance after shuffling barcodes among cells of the same library (200 shuffles). Values below 1 mean clonemates are closer.

**Table 4.**
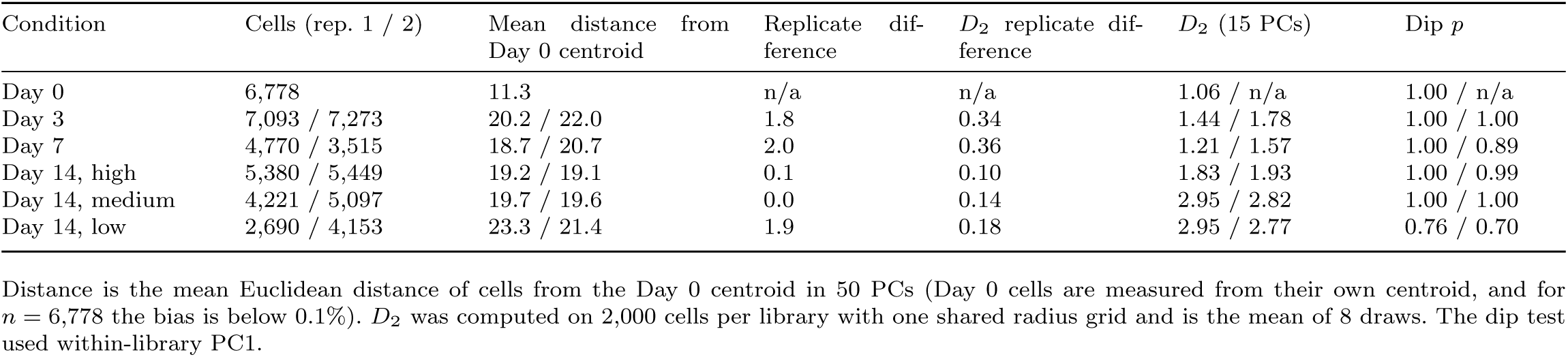
Per-library trajectory distance, correlation dimension and dip test in GSE150949.

| Condition | Cells (rep. 1 / 2) | Mean distance from Day 0 centroid | Replicate difference | *D*<sub>2</sub> replicate difference | *D*<sub>2</sub> (15 PCs) | Dip *p* |
|---|---|---|---|---|---|---|
| Day 0 | 6,778 | 11.3 | n/a | n/a | 1.06 / n/a | 1.00 / n/a |
| Day 3 | 7,093 / 7,273 | 20.2 / 22.0 | 1.8 | 0.34 | 1.44 / 1.78 | 1.00 / 1.00 |
| Day 7 | 4,770 / 3,515 | 18.7 / 20.7 | 2.0 | 0.36 | 1.21 / 1.57 | 1.00 / 0.89 |
| Day 14, high | 5,380 / 5,449 | 19.2 / 19.1 | 0.1 | 0.10 | 1.83 / 1.93 | 1.00 / 0.99 |
| Day 14, medium | 4,221 / 5,097 | 19.7 / 19.6 | 0.0 | 0.14 | 2.95 / 2.82 | 1.00 / 1.00 |
| Day 14, low | 2,690 / 4,153 | 23.3 / 21.4 | 1.9 | 0.18 | 2.95 / 2.77 | 0.76 / 0.70 |

Distance is the mean Euclidean distance of cells from the Day 0 centroid in 50 PCs (Day 0 cells are measured from their own centroid, and for *n* = 6,778 the bias is below 0.1%). *D*<sub>2</sub> was computed on 2,000 cells per library with one shared radius grid and is the mean of 8 draws. The dip test used within-library PC1.

One feature deserves a note. In both replicate pairs the Day 7 library lay closer to Day 0 than its Day 3 counterpart (replicate 1, 20.2 to 18.7; replicate 2, 22.0 to 20.7), a fall of 1.5 and 1.3. If the replicate 1 libraries and the replicate 2 libraries were cultured or processed together, the fall would be consistent across matched pairs, and a paired comparison could resolve it. The GEO metadata we used do not say whether they were, so we do not treat the pairs as matched. Whether the replicate difference reflects technical handling or biological drift between independent cultures does not change the unpaired reading. This does not show that no transition occurred between Day 3 and Day 7. What can be resolved is that cells moved away from the untreated profile by Day 3 and did not return within the 14 days sampled. Day 14 consists only of sorted cells and the earlier points are bulk, so sorting is a further difference between Day 14 and the rest.

### No library is bimodal, and the correlation dimension depends on the number of PCs

The dip test on within-library PC1 gave *p* `≥` 0.70 in all 11 libraries and *p* `≥` 0.95 in all five pooled replicate pairs (Table 4 and Additional file 3). There is no sign of two discrete modes in any condition, and pooling two replicates whose accuracy was 93% did not create one. The correlation dimension *D*_2_ at equal sample size (2,000 cells per library, one shared radius grid, 15 PCs) was 1.06 at Day 0, 1.44 and 1.78 at Day 3, 1.21 and 1.57 at Day 7, 1.83 and 1.93 for the high-retention Day 14 gate, 2.95 and 2.82 for the medium gate and 2.95 and 2.77 for the low gate. Replicate differences (0.10 to 0.36) were much smaller than differences between Day 0 and the Day 14 gates (0.8 to 1.9), so *D*_2_ distinguishes conditions here. It does not behave like the dimension of an attractor, though. In the Day 14 libraries it kept rising with the number of PCs (for example from 2.47 to 3.59 between 10 and 30 PCs in one low-retention library) without a plateau (Fig 3b), while Day 0 was nearly flat (0.94 to 1.06). We read *D*_2_ as a descriptive measure of how many directions the cells spread over and not as a property of a dynamical system.

### A within-library lineage comparison is immune to library effects

A comparison made inside one library is not affected by differences between libraries. GSE150949 records lineage barcodes for a large share of cells. Within every library (cells with a barcode shared by at least one other cell in the library, 509 to 3,461 cells and 90 to 1,000 clones), cells sharing a barcode were closer in 20-PC space than cells whose barcodes were shuffled within the same library. The ratio of mean squared distances was 0.76 to 0.89 (clonemates 11 to 24% closer), with permutation *p* = 0.005 (the smallest attainable with 200 shuffles) in all 11 libraries, including Day 0 (Fig 3c). This is a library-controlled result that is consistent with heritable expression differences between lineages, as in the original report [17]. It is not evidence of bistability. Heritable differences can arise from any slowly changing state, and the comparison has no information on stability or on dependence on path.

### Expression moves away from baseline in a single-library case study (GSE149383)

Raw expression matrices for GSE149383 [37] were merged, quality controlled and pooled (Methods), leaving 6,430 cells across eight conditions (Table 5). Sequencing depth initially drove the structure, since the first principal component (PC1) tracked log depth at *r* = 0.864. Regressing out log depth and mitochondrial fraction reduced this to *r* = −0.003. A linear correction cannot remove a non-linear or radial effect, however, and a depth gradient from the centre to the periphery of the embedding remains (Additional file 1), which we did not quantify. We therefore treat depth as a residual confounder and report depth-matched comparisons, a covariate-only classifier, count thinning and doublet detection (described in the next section). Downstream analyses use the depth-corrected embedding.

**Table 5.**
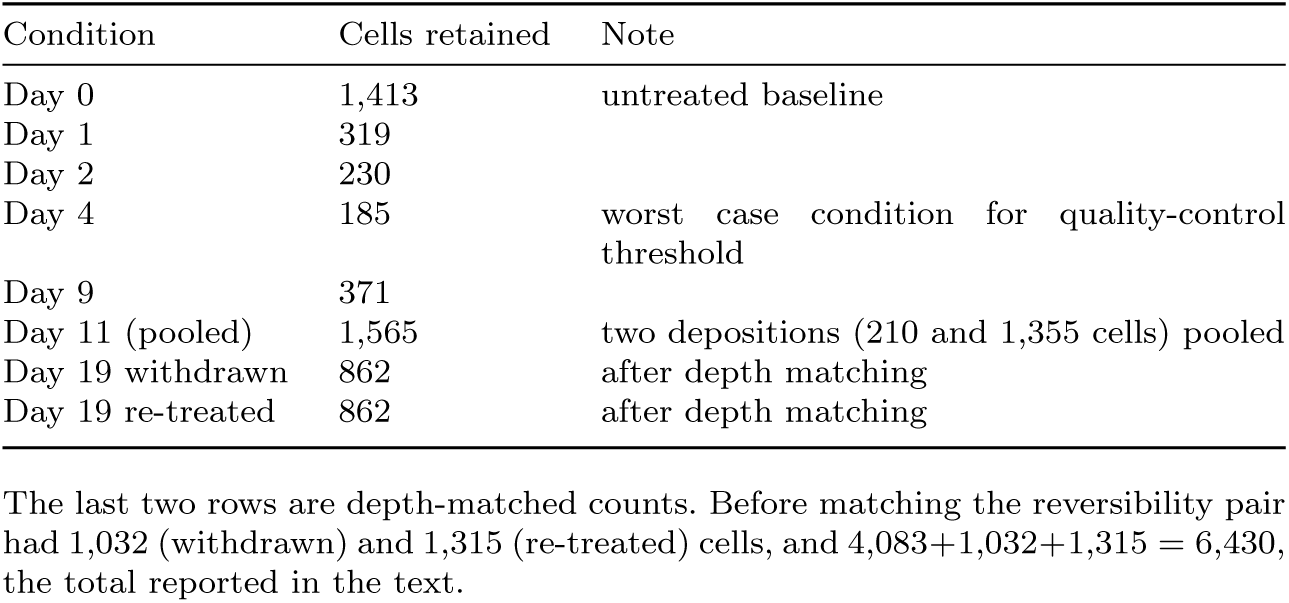
Per-condition cell counts entering the analysis after quality control and pooling.

| Condition | Cells retained | Note |
|---|---|---|
| Day 0 | 1,413 | untreated baseline |
| Day 1 | 319 |  |
| Day 2 | 230 |  |
| Day 4 | 185 | worst case condition for quality-control threshold |
| Day 9 | 371 |  |
| Day 11 (pooled) | 1,565 | two depositions (210 and 1,355 cells) pooled |
| Day 19 withdrawn | 862 | after depth matching |
| Day 19 re-treated | 862 | after depth matching |

The last two rows are depth-matched counts. Before matching the reversibility pair had 1,032 (withdrawn) and 1,315 (re-treated) cells, and 4,083+1,032+1,315 = 6,430, the total reported in the text.

We measured movement with diffusion pseudotime and the Euclidean distance from the Day 0 centroid. Mean pseudotime rose from 0.141 at Day 0 to 0.306 at Day 1 and then declined only partly (0.263, 0.274, 0.225 and 0.204 at Days 2, 4, 9 and pooled Day 11). Distance showed a similar overshoot with a later peak (9.53 at Day 0, 12.58, 12.78 and 13.19 at Days 1, 2 and 4, 11.92 at Day 9 and 11.85 at Day 11), settling at a plateau well above baseline (Fig 4a,b). Both measures derive from the same principal component analysis (PCA), so their agreement shows robustness to the choice of metric and not independence. Day 0 cells are measured from a centroid fitted on themselves, which biases their own distance downward. For *n* = 1,413 the squared distance changes by a factor of (1 − 1*/n*)*/*(1 + 1*/n*) = 0.9986, so this cannot explain the shift from 9.53 to 12.58. Day 11 pools two libraries (210 and 1,355 cells) that differ clearly (below), so that point is mostly one library. A cluster of 52 cells (0.8%) at extreme pseudotime, enriched for Day 0 and for high depth, was not a disconnected patch of the neighbour graph (14.0 mean neighbours of 15 requested), and removing it did not change the trajectory. The trajectory is compatible with an acute response followed by partial relaxation, but a cross-sectional design cannot separate transitions of individual cells from selective survival of pre-existing tolerant cells (Discussion). With one library per time point, the size of the partial return cannot be compared with replicate variation, which in GSE150949 was as large as the changes between later time points.

**Fig. 4.**
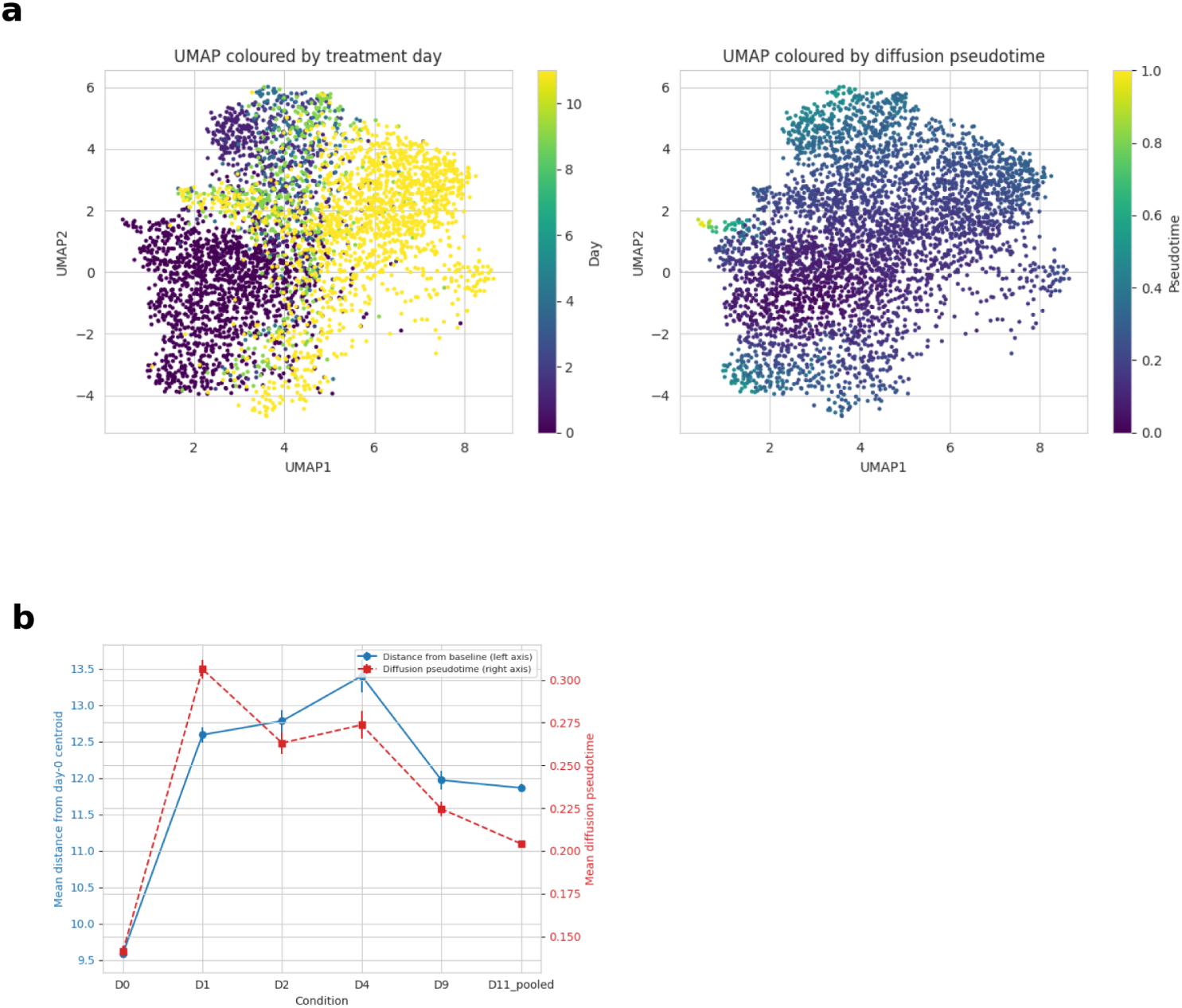
Expression moves away from baseline in the single-library erlotinib data. **a**, UMAP of the Day 0 to Day 11 erlotinib trajectory (GSE149383) coloured by treatment day (left) and diffusion pseudotime (right). **b**, Mean Euclidean distance from the Day 0 centroid (left axis) and mean diffusion pseudotime (right axis) across the six erlotinib timepoints. Both measures derive from the same PCA embedding, and Day 11 pools two libraries and is mostly one of them.

### Library effects are as large as the withdrawn versus re-treated difference

A logistic regression on the top 20 corrected PCs, with five-fold stratified cross-validation, separated depth-matched withdrawn and re-treated cells (862 each) with 88.5% accuracy (area under the receiver operating characteristic curve, AUC, 0.956, permutation *p* = 0.001, the smallest value 1,000 permutations allow, Table 6 and Fig 5a). Accuracy rose from 76% with 5 PCs to 89% with 50. A classifier given only log depth, mitochondrial fraction and genes detected reached 60% (AUC 0.61), so these covariates do not explain the separation. This shows that the two samples differ in expression, but not why.

**Fig. 5.**
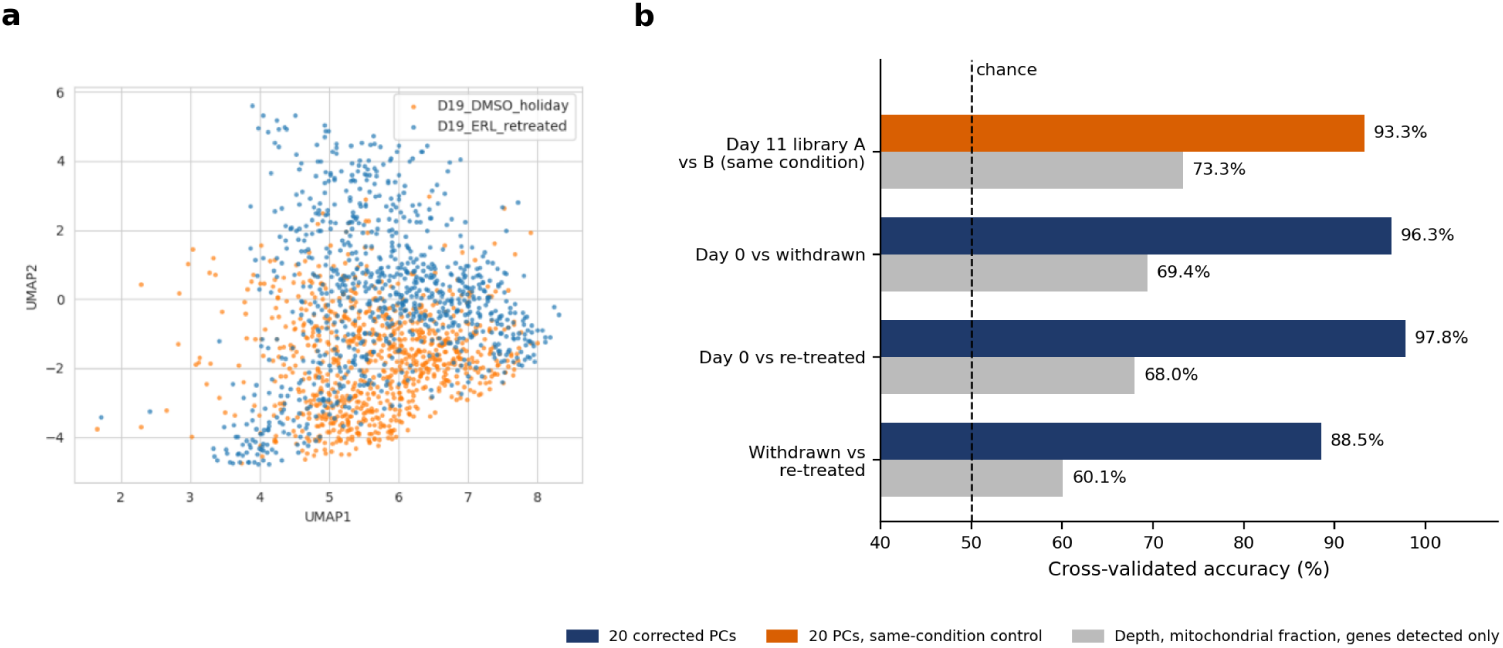
A library difference alone separates cells as well as the drug-history contrast. **a**, UMAP of the depth-matched withdrawn and re-treated cells (862 per group). **b**, Cross-validated accuracy of a logistic regression on 20 corrected PCs (blue and orange) and of a classifier given only log depth, mitochondrial fraction and genes detected (grey) for four depth-matched comparisons (values in Table 6). The orange bar is the control, two Day 11 libraries from the same condition, which separate better than withdrawn and re-treated cells. Dashed line, chance.

**Table 6.**
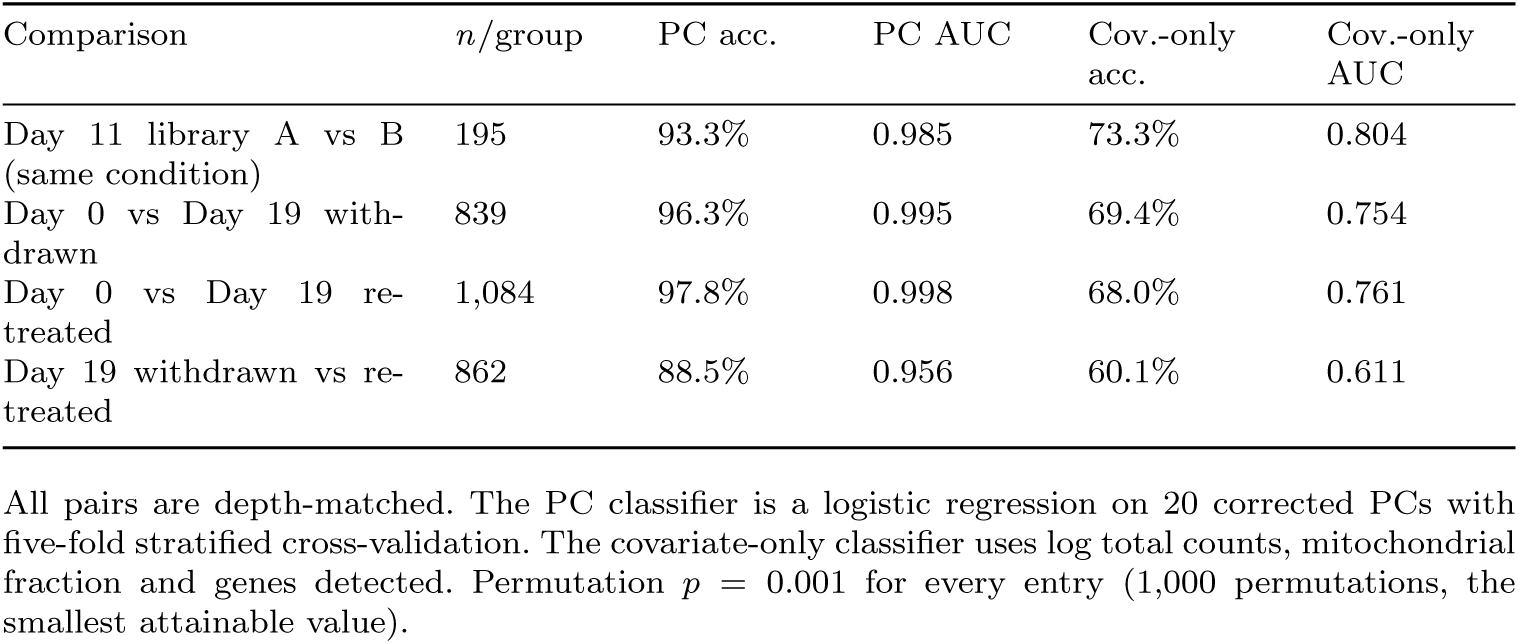
Library and depth controls.

The obvious control is to ask the same question of two libraries that should not differ. Day 11 was sequenced twice, in two independent depositions. Matched for depth at 195 cells per library, the same classifier separated them with 93.3% accuracy (AUC 0.985, permutation *p* = 0.001), better than it separated withdrawn from re-treated cells (Table 6 and Fig 5b). The covariate-only classifier reached only 60 to 73% through-out. The pattern held after binomial count thinning to a common depth and after removing cells that Scrublet called as doublets. Withdrawn versus re-treated stayed at 88.1% after either adjustment, and the two Day 11 libraries separated at 96.1% after thinning and 92.6% after doublet removal (Table 7). The 88.5% and 93.3% above are the values before these adjustments. Scrublet flagged 10% of re-treated cells and 10 to 16% of the two Day 11 libraries, against 0.2 to 0.3% of Day 0 and withdrawn cells, which is one more way in which the libraries differ. Differences between libraries in these data are therefore at least as large as the withdrawn versus re-treated difference, and depth does not account for them. The 88% cannot be attributed to drug history on this evidence, because a library difference alone could produce a similar value. The source of the library effect (sequencing run, culture or cell capture) cannot be deter-mined. The result also contradicts the check that originally justified pooling the two Day 11 libraries (*R*^2^ *<* 0.05 on each leading PC). That check examines one PC at a time and missed a difference that a 20-PC classifier finds easily.

**Table 7.** Depth and doublet controls.

| Check | Withdrawn vs re-treated | Day 11 library A vs B |
|---|---|---|
| After binomial thinning to 600 counts | 88.1% (AUC 0.952) | 96.1% (AUC 0.991) |
| After removing Scrublet doublets | 88.1% (AUC 0.953) | 92.6% (AUC 0.978) |

Accuracy and AUC are from the PC classifier (20 PCs). These adjustment checks used 200 permutations (*p* = 0.005 for every entry), whereas the unadjusted values in Table 6 used 1,000 (*p* = 0.001). The unadjusted depth-matched values (Table 6) are 88.5% for withdrawn versus re-treated and 93.3% for Day 11 library A versus B. The Day 11 contrast stays high, though not identical, after each adjustment.
The predicted doublet fraction was 0.2% for Day 0, 0.3% for withdrawn and 10.2% for re-treated cells, and 16.2% and 10.5% for the two Day 11 libraries. A residual radial depth gradient visible in Additional file 1 was not quantified.

Two features of this comparison need care. The replicate pair has 195 cells per library whereas the withdrawn versus re-treated pair has 862 per group, and accuracy rises with sample size. In simulation a replicate pair is separated at 85% with 50 cells per library, 91% with 100, 94% with 195, 96% with 400 and 96% with 862 (Additional file 2). Comparing at equal *n* would therefore, if anything, lower the 88.5% and widen the gap in favour of the replicate pair. We summarise the comparison as a replicate-referenced ratio *ρ* = (acc_contrast_−0.5)*/*(acc_replicate_−0.5). From the archived accuracies, *ρ* = (0.885 − 0.5)*/*(0.933 − 0.5) = 0.89. A value at or below 1 means the contrast is no more separable than two libraries that are supposed to be the same. Normalisation, depth regression and PCA were fitted on all cells before cross-validation. In simulation, fitting them inside the folds did not change accuracy (94.1% either way, Additional file 2).

The baseline comparison has the same limitation. Depth-matched Day 0 and withdrawn cells (839 each, median depth 1,608 versus 1,601 counts, Mann-Whitney *p* = 0.87) were separated with 96.3% accuracy (AUC 0.995), and Day 0 against re-treated cells (1,084 each) with 97.8% (AUC 0.998). These cells come from different libraries and differ by 19 days in culture, and the Day 11 control shows what a library difference alone can produce. The data therefore cannot show whether withdrawn cells moved back towards baseline, and they do not speak to the modest clinical activity of EGFR-TKI re-treatment [8].

### Exploratory analyses of the single-library erlotinib data are in Additional file 1

The remaining analyses of GSE149383 are exploratory and are reported in Additional file 1. At equal sample size and with one shared radius grid, the correlation dimension ordering reversed relative to a first pass at each condition’s own *n* (withdrawn 2.82, re-treated 3.32), it rose with the number of PCs without a plateau, and the log *C*(*r*) curves had no straight stretch. Convergent cross mapping between nine markers and a marker-free trajectory axis gave one signal that survived correction and surrogate tests (*EGFR*), but it sat at the 97.7th percentile of 300 expression-matched random genes (one-sided *p* = 0.027), the gene is flat across the time course, and transfer entropy did not support it. The permutation-entropy signal came from interpolated bins (13 of 30 series points), and no gene was significant on occupied bins alone. A comparison of nine markers between withdrawn and re-treated cells is subject to the library caveat above. We carry none of these forward as findings.

### Library-level permutation gives exact calibration: benchmark against alternative approaches

We compared *ρ_ED_* with the approaches a reader could use on the same data, in simulations with known truth (Methods; Additional file 3). Two conditions with *k* = 2, 3 or 4 libraries each were simulated with negative-binomial counts and 200 cells per library. There were five settings. In the first two the library effect was independent across genes or confined to five shared programs (low-rank). In the other three a second, tolerant state was occupied by 30% of cells (“state mix”), with an independent library effect, a low-rank library effect, or an independent effect plus variation in occupancy between libraries (standard deviation 10 percentage points; “varying”). Library effects were calibrated so that two libraries of one condition were separated at 87 to 94% accuracy, in the range of GSE150949. The difference between conditions was zero, a shift of the latent state (0.15, 0.30 or 0.60) or, in the state-mix settings, a change in occupancy of 0.10 or 0.20. Each setting had 200 simulations (150 for state mix), and every approach saw the same data (Fig 6 and Table 8).

**Fig. 6.**
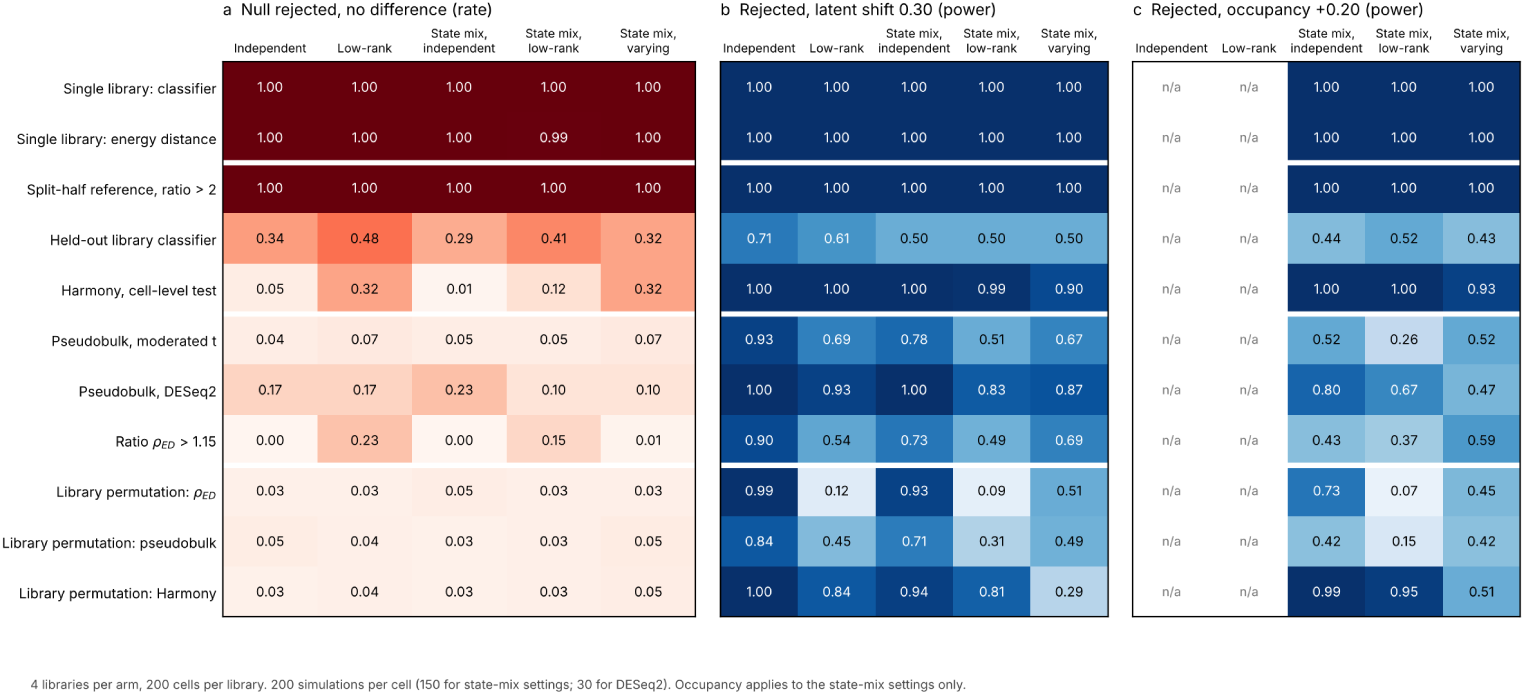
Benchmark against alternative approaches in simulations with known truth. **a**, Proportion of simulations with no difference in which each approach rejected the null (rule in Table 8). **b**, Rejection rate for a shift of the latent state of 0.30. **c**, Rejection rate for a change in tolerant-state occupancy of 0.20 (state-mix settings only). Four libraries per arm, 200 cells per library. Library permutation, exact test over the 35 splits of eight libraries. DESeq2: 30 simulations per setting.

**Table 8.**
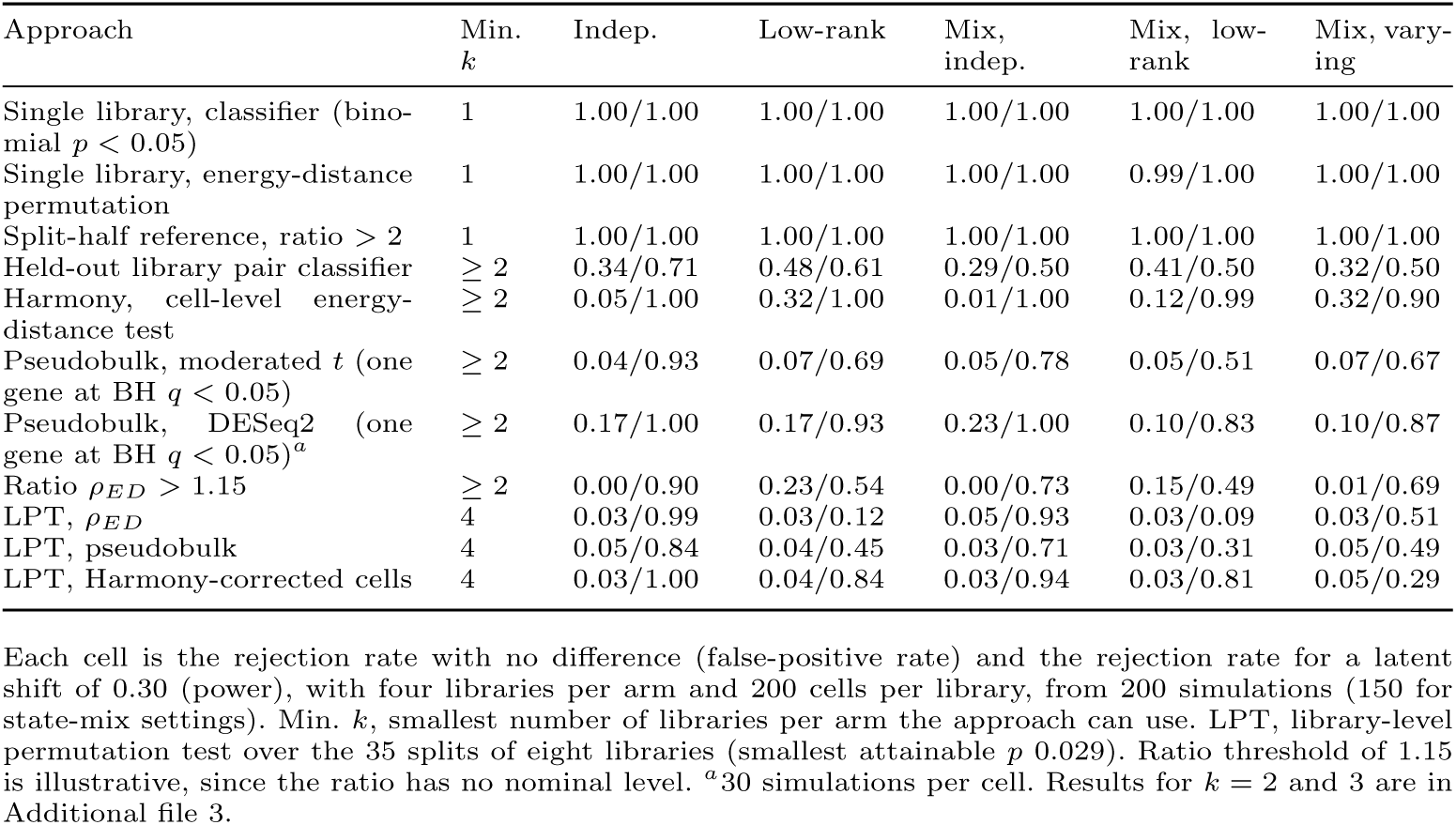
Null rejection and power of each approach at its nominal decision rule in simulation.

#### Single-library designs and a split-half reference measure the library effect

A cross-validated classifier and an energy-distance permutation test applied to one library per arm rejected the null in 99 to 100% of simulations with no difference, in all fifteen combinations of setting and *k*. A ratio of the between-condition distance to the split-half distance of a single library did the same. The classifier’s AUC for telling a shift of 0.30 from no shift was 0.45 to 0.56 across settings and *k*. In GSE150949 the split-half reference would have called every same-condition replicate pair different, with ratios of 8 to 156, and would have given ratios of 167 to 1,419 for the seven condition contrasts (Additional file 3). The accuracy ratio (acc_contrast_ − 0.5)*/*(acc_replicate_ − 0.5) saturated and was uninformative (AUC 0.48 to 0.55 with four libraries per arm). The replicate library is therefore the component that matters most in the reference.

#### Replicate-aware approaches with a parametric or cell-level null differ in validity

A classifier trained on one library per arm and tested on the held-out pair rejected the null in 25 to 48% of simulations, because cells within a library are not independent, and had low power. Harmony with library as batch, followed by a cell-level energy-distance test, was very sensitive (power 0.90 to 1.00 for a shift of 0.30 with four libraries per arm). It kept false positives at 1 to 5% with independent library effects but rejected the null in 8 to 40% of simulations with low-rank or varying effects (*k* = 2 to 4; 12 to 32% with four libraries). Pseudobulk testing with our moderated-*t* implementation rejected the null in 2 to 16% of simulations (4 to 7% with four libraries) and was more powerful than any exact test only when replicates varied in occupancy (0.67 against 0.29 to 0.51). DESeq2 on pseudobulk counts rejected the null in 7 to 23% of the simulations in which it was run (30 per setting, so these rates are imprecise). Augur, which pools cells from all libraries, separated simulations with a shift of 0.30 from simulations with none with an AUC of 0.70 to 1.00 at four libraries per arm in the settings tested, and gives no decision rule. A fixed threshold on *ρ_ED_* has no nominal level. A threshold of 1.15, near the null 95th percentile with independent effects, gave 0 to 1% false positives with independent effects and 15 to 23% with low-rank effects at *k* = 4 (up to 37% at *k* = 2), and a threshold of 2 gave at most 10% false positives but almost no power at these effect sizes.

#### A library-level permutation test is exact in every setting

For a statistic *T* that depends on the arm labels only through the split of libraries into two groups, the null hypothesis of no difference makes libraries exchangeable, so recomputing *T* for all 35 distinct splits of eight libraries gives an exact *p* value whatever the structure of the library effect (Methods). The statistic may include steps fitted to library identity, such as Harmony. LPT held false positives at 3 to 5% in all five settings for *ρ_ED_*, pseudobulk and Harmony-corrected cells, as designed. Power differed with the statistic (Fig 6b,c). For a latent shift of 0.30, LPT on Harmony-corrected cells reached 1.00, 0.84, 0.94 and 0.81 in the independent, low-rank, state-mix independent and state-mix low-rank settings, against 0.99, 0.12, 0.93 and 0.09 for LPT on *ρ_ED_* and 0.84, 0.45, 0.71 and 0.31 for LPT on pseudobulk. For an occupancy change of 0.20 the powers were 0.99, 0.95 and 0.51 (Harmony), 0.73, 0.07 and 0.45 (*ρ_ED_*) and 0.42, 0.15 and 0.42 (pseudobulk). When replicates varied in occupancy, LPT on Harmony-corrected cells was the least powerful of the three for a latent shift (0.29 against 0.51 and 0.49). The smallest attainable *p* is 0.029 with four libraries per arm, 0.10 with three and 0.33 with two, so no library-level test can reject at *p <* 0.05 with fewer than four.

#### Two pre-specified variants of the ratio did not help, and the ablations show where it is stable

We pre-specified two label-free variants of *ρ_ED_* that use the replicate libraries to estimate library noise: inverse-variance weighting of the PCs (whitened) and removal of the leading directions of library variation (projected). Projection removed the signal (AUC 0.06, 0.16 and 0.24 in three settings at *k* = 4). Whitening raised the AUC with low-rank effects (0.85 against 0.78) but lowered it with state mix and varying occupancy (0.84 against 0.95), and its LPT power was lower in the independent setting (0.89 against 0.99). Neither is recommended. Replacing the replicate reference by split halves or the energy distance by a classifier accuracy reduced the AUC to 0.48 to 0.66, whereas the centroid distance performed as well as the energy distance (Fig 7c). With two libraries per arm and independent effects the AUC of *ρ_ED_* was 0.50, 0.95, 0.97 and 0.98 with 5, 10, 20 and 50 PCs and 0.91, 0.97 and 0.98 with 100, 200 and 400 cells per library, and with low-rank effects it was flat across PCs and varied by at most 0.06 across cells per library (Fig 7a,b).

**Fig. 7.**
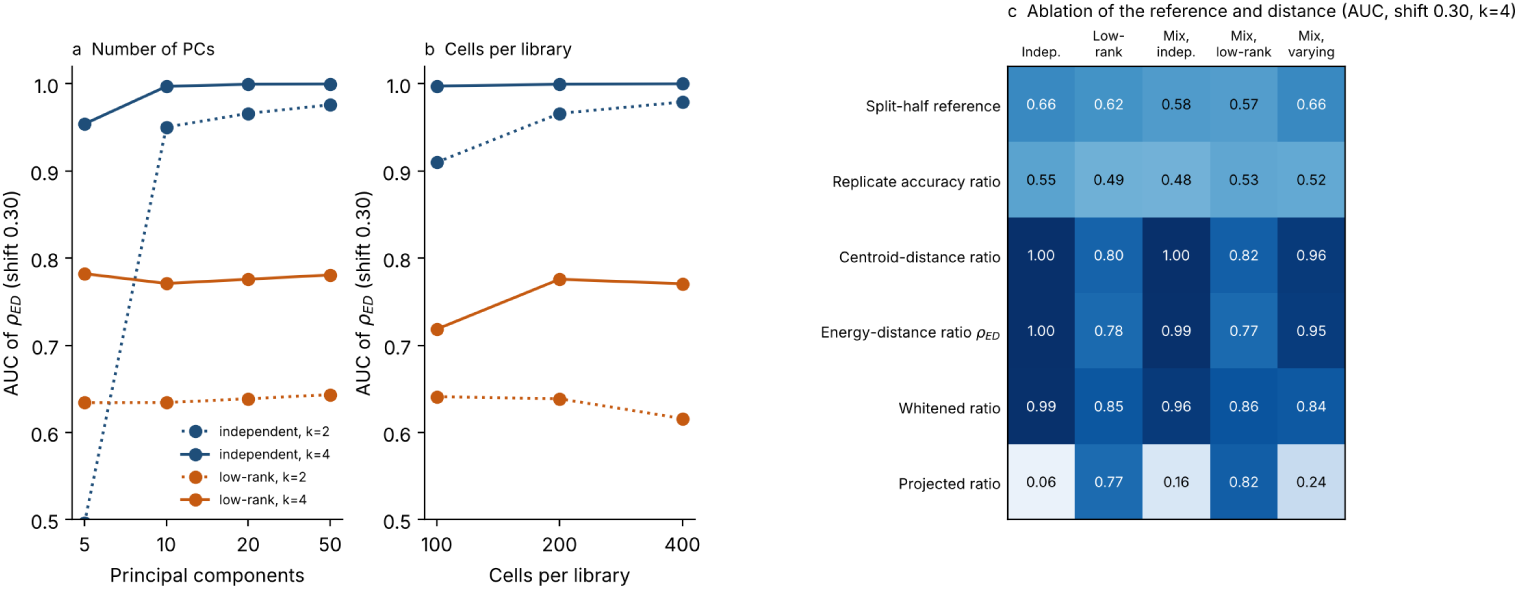
Ablations of the replicate-referenced ratio. **a**, AUC of *ρ_ED_* for a shift of 0.30 against the number of PCs. **b**, AUC against cells per library (20 PCs). Colour shows the library-effect structure and line style shows libraries per arm *k*. **c**, AUC of variants of the ratio at *k* = 4: split-half reference, replicate accuracy ratio, centroid-distance ratio, energy-distance ratio *ρ_ED_*, and the whitened and projected variants.

These results do not show that one approach is best in every setting. They show that a replicate library is required as the reference, that exact library-level permutation is the only calibration that held in all five settings, that the statistic inside it determines power, and that Harmony-corrected cells gave the most power of the exact tests in most settings. With fewer than four libraries per arm, no exact test exists, and *ρ_ED_* remains a replicate-referenced effect size. It is the quantity reported for the real datasets below.

### Simulations of the toggle switch show when the other checks work and when they do not

Real data cannot say what an analysis would return when the truth is known. We therefore ran simulations of a two-gene toggle switch observed through negative-binomial counts, with library effects calibrated to a replicate accuracy of 93% (Additional file 2). Under five noise structures a single-library contrast was called separable in every simulation, including those with no biological difference, and a replicate pair separated at only 56% already gave 77% false positives. The replicate-referenced distance ratio kept its false-positive rate and detected large effects when the library effect was high-dimensional. It needed three or four libraries per arm for small effects (power for a small monostable shift rose from 20% with two libraries per arm to 63% with four) and had a heavy-tailed null when the library effect was low-dimensional and only two libraries per arm were available. A dip test run separately in each library and required to agree gave no false positives, whereas pooling two libraries made every simulated monostable population look bimodal. Correlation dimension followed the number of PCs and the sequencing depth and not the latent dimension, and permutation entropy of interpolated series was significant for 90 to 95% of genes unrelated to the trajectory. The simulations are simple, and in the real replicate data pooling replicates did not make the dip test fire, so their numbers calibrate the argument and are not predictions for any dataset.

## Discussion

GSE150949 shows the framework at work, because its replicate libraries supply the scale that single-library datasets lack. Halves of one library were indistinguishable, replicate libraries of one condition were separable at 59 to 93%, and the contrasts of main biological interest, time in drug and sort gate, exceeded the replicate distance 4.5-to 51-fold at 20 PCs (at least 3.7-fold from 10 to 50 PCs, Table 3). After osimertinib, PC9 cells therefore moved away from their untreated expression profile by Day 3 and did not return within 14 days. This is the most secure result of the study. Finer features are a different matter. The fall in distance from Day 0 between Day 3 and Day 7 was smaller than the replicate differences, so an overshoot and partial relaxation along that axis, or any rise, fall and rise shape, cannot be read from one library per time point, even though the Day 3 and Day 7 populations differ. A distance ratio expresses the comparison directly and, unlike classifier accuracy, it does not saturate when the replicate effect is large. It is a diagnostic and not a test. With two libraries per arm it has no library-level null, its value depends on the number of PCs, and it can be inflated by chance when the replicate difference is concentrated in few directions. No library showed two modes, and *D*_2_ separated conditions but kept rising with the number of PCs at Day 14. The one comparison in these data that library effects cannot confound is the lineage comparison inside a library, in which clonemates were closer than shuffled cells in every library. Comparisons of this kind are not possible in single-library designs unless lineage barcodes or hashing are available.

The benchmark shows why the reference must be a replicate library and how to test against it. Single-library tests and a split-half reference rejected the null in nearly every simulation with no difference. Replicate-aware approaches with a cell-level or parametric null were valid in some settings and not in others. Harmony followed by a cell-level test was the most sensitive but rejected the null in up to 40% of simulations with low-rank or varying library effects, and pseudobulk testing was valid with four libraries per arm but not with two in the low-rank setting. A library-level permutation test was exact in all five settings, so we recommend it whenever four or more libraries per arm are available, with Harmony-corrected cells as the default statistic and *ρ_ED_*reported as the effect size. When replicates vary in the occupancy of the tolerant state, LPT on Harmony-corrected cells lost power for a latent shift, and LPT on *ρ_ED_* or on pseudobulk should be examined alongside it. With two or three libraries per arm, as in the datasets analysed here, no exact test exists. In that regime *ρ_ED_*with replicate pairs, a held-out replicate, and lineage barcodes give a scale and a check, not a test.

The central control is also the simplest. Two Day 11 libraries, with the same drug and the same day, were separated with 93% accuracy, more than the 88% reached for withdrawn against re-treated cells. A classifier result in this dataset can therefore reflect library identity at least as much as treatment. The two arms also differ in what they received last (eight drug-free days versus six drug-free days and two days back on erlotinib [37]), so the comparison was never a clean test of drug history. Depth, mitochondrial fraction, count thinning and doublets do not explain the library effect, and we cannot say whether it comes from the sequencing run, the culture or cell capture. It follows that the Day 0 versus Day 19 accuracies (96 to 98%) do not tell us whether withdrawn cells went back towards baseline, and that the pooled Day 11 point, 87% of which is one library, is less solid than it looks. The range of replicate separability in GSE150949 (59 to 93%) shows that a value of 93% between two libraries of one condition lies within what a replicated 10x experiment can produce. The platforms differ, so this is a statement about plausibility and not a comparison of the datasets. The general point is that in a single-library design a replicate effect is likely present but cannot be measured.

The correlation dimension did not hold up either. At equal sample size in the erlotinib data the ordering flipped, and even then the value depends on the number of PCs, has no plateau, and the log *C*(*r*) curves bend throughout. *D*_2_ also rose with the number of PCs in the Day 14 libraries of GSE150949, and a simulation with a known latent dimension of 2 made the same point (Additional file 2). We do not think these data allow *D*_2_ to be read as a measure of attractor geometry, and the dip test and mixture model give no sign of two modes. The cross-mapping, transfer-entropy and permutation-entropy analyses rest on 30-point series of cross-sectional means with 13 interpolated points, and the one gene that stood out (*EGFR*) did not survive comparison with expression-matched random genes and other checks. We classify it as an exploratory observation and do not offer it as a candidate for perturbation experiments (Additional file 1).

GSE150949 has limits too. It has two replicates per condition but only one Day 0 library, Day 14 is sorted while earlier points are bulk, and the replicate cultures differ in biology as well as in handling, so the replicate effect is an upper bound on technical variation. Using it as the reference does not penalise genuine heterogeneity. It asks only whether a between-condition difference exceeds the difference between independent cultures of one condition, which is the variation any claimed state change must stand out from whatever its origin. A reference that was purely technical would be smaller and would make more contrasts look significant, but it would not be avail-able from these data. The intervals in Tables 2 and 4 reflect subsampling of cells and not uncertainty about the libraries, which with two per arm is large. Whether replicate effects in other persister datasets resemble these is not known, and the 10x platform differs from Drop-seq. We also cannot say anything about osimertinib combined with chemotherapy [38, 39], because no public single-cell dataset with that arm was available to us.

What a better design buys can be seen from the replicate data and from the simulations (Additional file 2). Two libraries per condition allow a replicate-referenced contrast. In simulation, three or four per arm were needed to detect small effects, and the distance ratio had a heavy-tailed null when the library effect was low-dimensional, so a permutation test across libraries is advisable, and it can reach *p <* 0.05 only with four or more libraries per arm. Hashed libraries [40] and lineage barcodes allow comparisons inside a library. We do not propose a new estimator of attractor structure from snapshots. What behaved correctly was narrower and consisted of replicate-referenced contrasts, a dip test required to agree across libraries, and within-library lineage comparisons. The benchmark supports the first of these directly and the other two through the real replicate data and the simulations in Additional file 2.

### What these results do and do not show

Two statements are easy to run together, and we want to keep them apart. We do not claim chaos, which would need a positive Lyapunov exponent from a long, densely sampled series, and we avoid the terms bistability and hysteresis, which require two stable states under one shared condition and a demonstrated dependence on path. The two arms of GSE149383 also differ in final drug exposure, so some difference between them is expected and would not by itself indicate bistability. The first statement is that these data do not support attractor-style inference. We think the controls above establish this for the datasets analysed. The second is that the cancer-attractor hypothesis is wrong, or that drug-tolerant states are not bistable. These data cannot say that, and we do not claim it. A design with replicate libraries for every condition, and with dense longitudinal sampling or lineage tracing, could reveal dynamical structure that a single-library, six-timepoint dataset cannot reach. Our negative result applies to these two public datasets and, with caution, to similar single-library designs, and to the estimators we tested on them. It does not apply to the biology. The estimators themselves are not necessarily at fault, since correlation dimension and CCM were developed for long, densely sampled series and we applied them in a regime they were not built for. Table 9 lists the controls we consider useful and what each showed here.

**Table 9.**
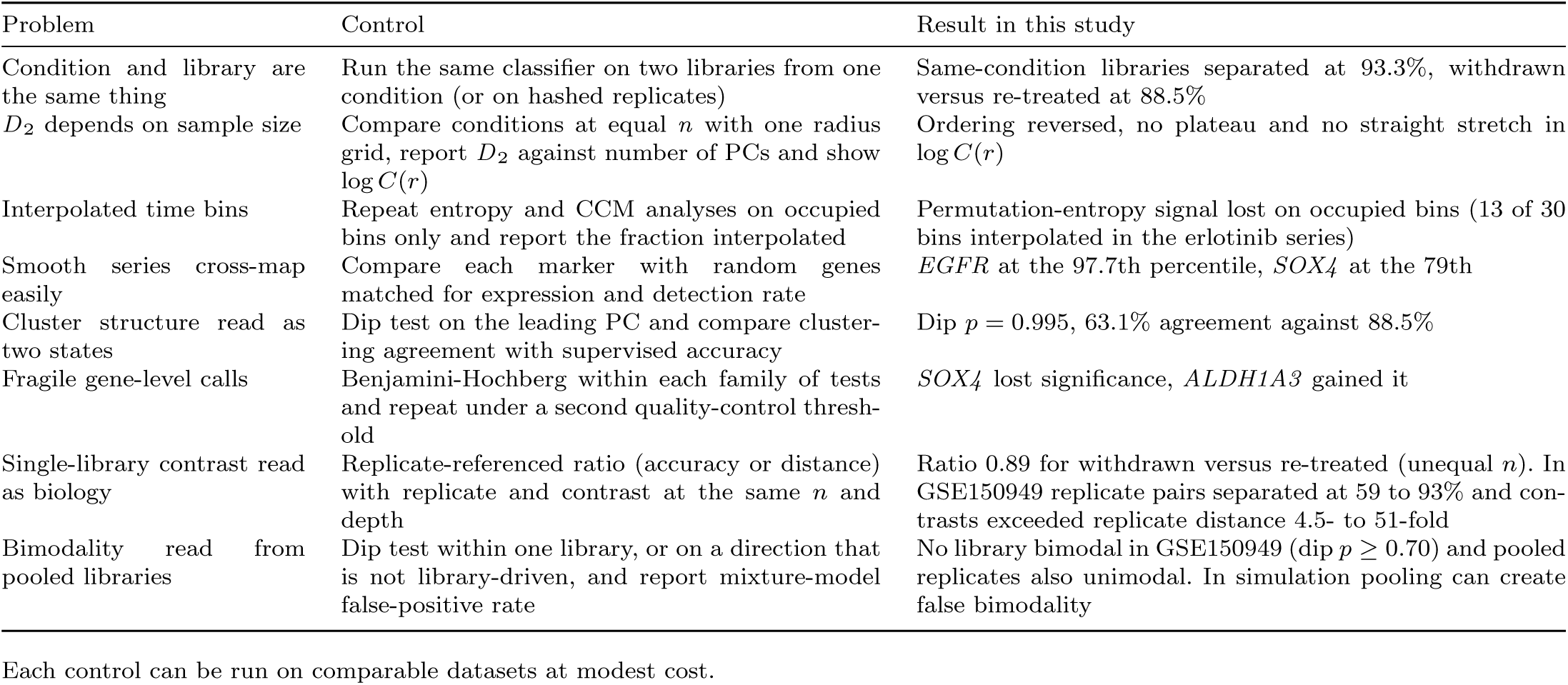
Controls for each problem and what they showed here.

### Scope and design guidance

#### What the two datasets can support

The designs are cross-sectional. No cell is observed at more than one timepoint, so trajectories are population-level inferences, and because the drug kills sensitive cells throughout, a trajectory may reflect selection of tolerant cells as well as transitions of individual cells. GSE150949 has two libraries per condition, one Day 0 library and sorted cells at Day 14, and its replicate cultures differ in biology as well as technique. The GEO metadata do not say whether replicate libraries are matched across time points or how they were cultured and processed, so the biological and technical sources of the replicate effect were not separated and pairing could not be used. The intervals in Tables 2 and 4 reflect subsampling of cells and not uncertainty about the libraries. The main analysis used 20 PCs and 1,300 cells per group. The distance ratio was recomputed at 10 and 50 PCs with 800 cells per group (Table 3), whereas classifier accuracy and the other statistics were computed at 20 PCs. GSE149383 has one library per condition. Its Drop-seq depth is shallow (median about 1,300 to 2,000 UMIs in the reversibility arm, with 300 as the quality-control minimum), a radial depth gradient remains after linear correction (Additional file 1), and no lane or batch identifiers are available, so the source of its library effect (sequencing run, culture or cell capture) cannot be assigned. Its second quality-control threshold shares about 90% of its cells with the first and is a sensitivity check and not independent confirmation. The replicate-referenced ratio of 0.89 for GSE149383 uses accuracies computed at unequal sample sizes and has no interval. Both datasets come from one cell line on different platforms, so the findings describe these data.

#### Designs that would extend the result

Four or more libraries per arm make an exact library-level test possible. Lane and batch identifiers, hashed libraries [40] and lineage barcodes would let the source of the replicate effect be separated from the biology. Public datasets with replicate libraries or hashed samples from other systems, for example cell-line mixtures profiled in replicate [41] or multiplexed donors [42], are the natural next test of the benchmark on real libraries with known truth, and we did not analyse them here. Whether replicate effects of the size seen in GSE150949 occur in other persister systems is not known.

#### Scope of the benchmark

The simulations (Additional file 2) use a two-gene circuit and simple technical-noise models, with library effects calibrated to one observable and shifts expressed in latent units, so their numbers calibrate the argument and are not predictions. Library-level permutation assumes that libraries are exchangeable under no difference, which holds in the simulations (equal numbers of cells, the same processing) and may not hold when libraries differ systematically in size, chemistry or time of processing. Its exactness needs four or more libraries per arm, and the two real datasets here have two or one, so it could not be applied to them. We implemented the moderated pseudobulk test ourselves, ran DESeq2 through pydeseq2 and Augur through pertpy on 30 simulations per setting, and used Harmony at its default set-tings. kBET, mixed models and neighbourhood-based differential abundance were not benchmarked, because R was not available to us, and are discussed only conceptually. Effect sizes in the simulations (median *ρ_ED_* 1.2 to 1.9) are smaller than the contrasts in GSE150949 (4.5 to 51), and the 1.15 and 2 thresholds are illustrative.

#### Terminology

“Coupling” means statistical coupling and not causation. We make no claim of bistability, hysteresis, a vector field, chaos or a fractal attractor.

## Conclusions

Contrasts in cross-sectional single-cell data should be judged against the difference between replicate libraries of one condition. In simulations with known truth, single-library tests and a split-half reference rejected the null in nearly every run with no difference, and Harmony followed by a cell-level test did so in up to 40% of runs with low-rank or varying library effects. A library-level permutation test was exact in all five settings with four libraries per arm, and applying it to Harmony-corrected cells gave the most power of the exact tests in most settings. With fewer libraries, *ρ_ED_* remains a replicate-referenced effect size. In a public 10x dataset with replicate libraries, halves of one library were indistinguishable, replicate libraries of one condition were separable at 59 to 93%, and contrasts between times in drug and sort gates exceeded the replicate distance 4.5- to 51-fold at 20 PCs (at least 3.7-fold from 10 to 50). Cells moved away from their untreated profile by Day 3 and did not return in 14 days, and clonemates were closer than random cells in every library. The fall in distance from Day 0 between Day 3 and Day 7 was smaller than replicate differences, although the two populations differ. No library was bimodal, and the correlation dimension depended on the number of principal components. In the Drop-seq dataset with one library per condition, two libraries of the same condition were separated more readily (93%) than withdrawn and re-treated cells (88%), so classifier accuracy there cannot be attributed to drug history, and the correlation dimension, permutation entropy and cross-mapping analyses did not hold up. Attractor language would be justified by designs that can show stability and path dependence, such as four or more replicate or hashed libraries for every condition, lineage barcoding in a withdrawal and re-treatment experiment [16, 17] and densely sampled longitudinal data. The code supplied with this paper runs these checks on comparable datasets.

## Methods

### Cell lines and ethics

We did no new experiments. The PC9 cells (an EGFR exon-19-deletion line) were grown and sequenced by the authors of the original studies [17, 37], and we refer to those papers for the source and authentication of the line. We did not obtain, culture or handle cells. No human participants or vertebrate animals were involved, so no ethics approval was needed.

### Datasets and preprocessing

Raw digital gene expression matrices for GSE149383 were obtained from the NCBI Gene Expression Omnibus [37]. The data were generated by Drop-seq [43]. We used the nine PC9 samples of the 2 *µ*M erlotinib time course. These were Day 0 (untreated), Day 1, 2, 4 and 9, two independently sequenced Day 11 depositions, and the reversibility arm (11 days of erlotinib followed by 8 drug-free days, “withdrawn”, DMSO, or 6 drug-free days and 2 days of re-exposure, “re-treated”), following the original study [37]. Two further depositions labelled as Day 0 and Day 2 replicates were excluded because MD5 hashing and cell-barcode-set comparison showed they were byte-for-byte duplicate re-uploads of other GEO samples. The two independent Day 11 depositions were pooled after a check on the leading PCs (*R*^2^ *<* 0.05 on each), which later proved insensitive (Results). Because per-sample files omit genes with zero counts, the nine retained files were merged on the union of observed gene symbols (30,297) with implicit zeros, giving 31,937 raw cells. Twenty-nine cells with zero total UMI counts were removed. The raw per-cell count distribution was strongly right-skewed (mean 653.7, median 93.0, s.d. 2,074), which is consistent with a Drop-seq library in which most droplets contain ambient RNA. Because depth varied substantially across conditions, per-condition barcode-rank (“knee”) plots were used to locate the transition between cells and background. Cells were kept if total UMI count was at least 300, detected genes at least 30 and mitochondrial fraction below 30%, a threshold chosen so that every condition retained at least 185 cells (worst case Day 4). Quality control was performed before normalisation. This left 6,430 cells across eight conditions (Table 5).

Counts were rescaled to 10^4^ per cell and log-transformed,

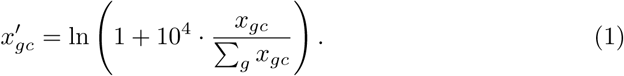

Genes were ranked by normalised dispersion within expression-level bins, and the top 2,000 were retained together with *EGFR* and eight literature drug-tolerance markers (*TACSTD2*, *SERPINE1*, *CD24*, *MET*, *ALDH1A3*, *SOX4*, *GPRC5A*, *CYP1B1*), which were force-included irrespective of dispersion rank (*p* = 2,009 genes). The matrix was column-standardised and decomposed by principal component analysis [44]. Because PC1 of the naively normalised data correlated with log sequencing depth (*r* = 0.864), we fit an ordinary least squares regression per gene,

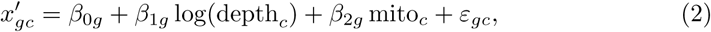

and kept the residuals, equivalent to scanpy’s regress out [45]. Depth varied continuously within every condition rather than being tied to a discrete batch label, for which a categorical correction such as Harmony would be appropriate [30], so a per-gene linear regression was used. It reduced the correlation between PC1 and depth to *r* = −0.003. Fifty components were retained for the neighbour graph and diffusion pseudotime, the top 15 for *D*_2_ (to limit the downward bias of the estimator at high embedding dimension) and the top 20 for classification. Sensitivity to these choices is reported in the code repository.

### Trajectory measures

Diffusion pseudotime was computed from a *k* = 15 nearest-neighbour graph in the corrected 50-PC space, following Haghverdi and colleagues [46]. Pairwise distances are converted to a Gaussian affinity kernel and row-normalised into a Markov transition matrix, and pseudotime is the diffusion distance from a root cell, here the cell closest to the Day 0 centroid, to every other cell. As an assumption-light cross-check requiring no root-cell choice, we also computed the Euclidean distance of every cell from the Day 0 centroid in the corrected PC space, dist(*c*) = ||*x_c_* − *µ*_Day 0_||_2_. Because both measures derive from the same PCA, their agreement shows robustness to the choice of metric and not independence. Euclidean distance was adopted as the primary trajectory measure.

### Correlation dimension

For *N* points (cells in the top 15 corrected PCs) the correlation sum is *C*(*r*) = 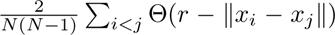, the fraction of distinct cell closer than *r* [47] Where *C*(*r*) ∝ *r^D^*^2^, *D*_2_ is the slope of ln *C*(*r*) against ln *r* over a scaling window (15 to 60% of a logarithmic grid spanning the 5th to 95th percentile of pairwise distances). *D*_2_ is a descriptive measure and not evidence of a fractal or chaotic attractor. The estimator is biased downward for small *N* and for embedding dimensions above about five [48], so values are comparable only at equal *N* within this study. The first-pass analysis used 200 bootstrap resamples (80% of cells, with replacement, zero-distance pairs removed) at each condition’s own *n*. The recomputation draws subsamples with-out replacement at a common *n* (185 cells, the smallest condition), with one radius grid shared by all conditions and *i* `̸`= *j* pairs. The difference *D*_2_(withdrawn)−*D*_2_(re-treated) was tested at equal *n* with a label-permutation test. *D*_2_ was recomputed with 10, 15, 20 and 30 PCs (Additional file 1).

### GSE150949 replicate analysis

The genes-by-cells count matrix (22,166 genes, 56,419 cells) and the cell metadata (library identifier, time point, sort gate, mitochondrial fraction, lineage barcode) were taken from the series’ GEO supplementary files [17]. All cells had passed the original filters (at least 1,000 genes, at least 1,859 UMIs, mitochondrial fraction at most 0.10). Counts were scaled to 10^4^ per cell and log-transformed. The 2,000 most variable genes were chosen by dispersion normalised within 20 bins of mean expression (genes with mean log expression 0.0125 to 3). Log UMI count and mitochondrial fraction were regressed out, genes were standardised, and 50 principal components were computed by randomised singular value decomposition (PC1 correlated with log depth at *r* = 0.000). The matrix was streamed in chunks of 250 genes to fit in 3 GB of memory. Library identity was read from the sample name column (`0, 3_rep1, 3_rep2, 7_rep1, 7_rep2, 14_rep1` and `14_rep2` with high, medium and low retention of the reporter, where low retention means more divisions [17]).

For each draw, 1,300 cells per library were sampled without replacement. Split-half floors compared two random halves of one library (2,600 cells). Replicate comparisons used the two libraries of one condition. Classification used the logistic regression on 20 PCs described below. Energy distance between point clouds *A* and *B* is 2E||*A* − *B* || − E ||*A* − *A^′^* || − E ||*B* − *B^′^* || [32]. For a contrast between arms with two libraries each, *ρ_ED_* is the mean energy distance over the four library pairs across arms divided by the mean over the two replicate pairs within arms. The accuracy ratio is *ρ* = (acc_contrast_ − 0.5)*/*(acc_replicate_ − 0.5) with the accuracy of the first library of each arm and the mean replicate accuracy of the replicated arms. Held-out replicate accuracy was obtained by training on one library of each arm and testing on the other, and the reverse, averaged. For contrasts with Day 0 the Day 0 library was split into halves for training and testing. The replicate difference direction was the difference between replicate centroids in 20 PCs. The dip test used PC1 of a PCA fitted within each library (or within the two pooled replicates) on the standardised 2,000-gene matrix. The correlation dimension used 2,000 cells per library (8 draws), 10, 15, 20 or 30 global PCs and one radius grid per number of PCs, built from a random sample of 2,000 cells of the whole dataset, with the slope fitted over 15 to 60% of a logarithmic grid between the 5th and 95th percentile of pairwise distances. Trajectory distances were Euclidean distances in 50 PCs from the Day 0 centroid. For lineage concordance, cells of a library with a recorded lineage barcode shared by at least one other cell of that library were used. The statistic was the mean squared 20-PC distance among cells sharing a barcode, compared with the mean of 200 permutations that shuffled barcodes among those cells, with *p* = (*r*+1)*/*201. Intervals are 2.5 to 97.5% ranges over 20 random draws of cells (the same draws throughout, with fixed seeds). The analysis code is available from the repositories listed under Availability of data and materials. 3

### Library distance matrix and sensitivity to the number of PCs

Using the processed 50 PCs of all 56,419 cells, 800 cells per library were drawn without replacement (5 draws, fixed seed). Energy distance was computed between every pair of libraries and, on the diagonal, between two random halves of each library, with unbiased within-sample terms, using the first 10, 20 or 50 PCs. The ratio for a contrast is the mean distance over all library pairs across the two arms divided by the mean over the replicate pairs within the replicated arms.

### Benchmark against alternative approaches

The simulation is in method comparison.py and the driver in run benchmark v2.py (Additional file 3). Each simulation draws 2*k* libraries of 200 cells (100 or 400 in the sample-size ablation), *k* per arm, over *G* = 300 genes. Gene log-means are *b_g_*∼ *N* (−5, 1) plus 0.5 times a two-dimensional latent state *z* ∼ *N* (0, 0.5^2^*I*) multiplied by loadings (40% of genes informative, loadings *N* (0, 1)). Arm B has its latent state shifted by (*δ,* −*δ*) with *δ* ∈ {0, 0.15, 0.30, 0.60}. In the state-mix settings a fraction *f* = 0.3 of cells in every library has its latent state moved by (1.2, −1.2) (a second, tolerant state), arm B has *f* raised by 0.10 or 0.20 in the occupancy experiments, and in the varying setting *f* is drawn per library from *N* (*f,* 0.10^2^). A library effect adds a gene-wise shift to the log-mean, either independent, *N* (0*, τ* ^2^), or low-rank, a random combination of five shared programs plus independent noise with standard deviation 0.02. The scales were chosen on a grid so that two libraries of one condition were separated at 87 to 94% by the 20-PC logistic regression (mean over the simulations of each setting): *τ* = 0.18 (independent, with and without state mix), 0.09 (low-rank, standard deviation of the program weights), and *τ* = 0.16 with varying occupancy. Counts are negative binomial with variance *µ* + 0.5*µ*^2^ and median depth 1,500 UMIs with log-normal variation (SD 0.5). Counts are scaled to 10^4^, log-transformed, standardised and decomposed by one PCA over all libraries. Seeds are fixed. Single-library approaches use library 1 of each arm. The classifier is a logistic regression with five-fold cross-validation and a one-sided binomial test, and the distance test is a label-permutation test of the energy distance (199 permutations). The held-out classifier is trained on *k*−1 libraries per arm and tested on the left-out library pair, with a binomial test on pooled cells. Harmony [30] (harmonypy 2.1.0, default settings, library as batch) is applied to 20 PCs and followed by the cell-level energy-distance permutation test between arms. Pseudobulk testing takes the mean log-normalised expression per library and gene, computes a two-group *t* statistic across libraries, and moderates the variances with the empirical-Bayes procedure of Smyth (equal residual degrees of freedom, estimated prior degrees of freedom). It calls a difference when one gene has a Benjamini-Hochberg *q <* 0.05. DESeq2 (pydeseq2 0.6.0, design ∼ condition, Wald test) was run on counts summed per library and Augur (pertpy 1.5.1, random forest, 10 subsamples of 100 cells) on all cells, in 30 simulations per setting for *k* = 2 and 4 (`external_tools.py`). *ρ_ED_*is the mean energy distance over library pairs across arms divided by the mean over replicate pairs within arms (20 PCs). Ablations replace the replicate reference by the mean split-half energy distance of single libraries, the energy distance by the Euclidean distance between library centroids or by the accuracy ratio, and use 5, 10, 20 or 50 PCs. The whitened variant weights each PC by 1*/*(*v_j_* + 0.1*v̅*), where *v_j_* is the variance of the library centroids’ deviations from their grand mean, and the projected variant removes the leading 2*k* − 3 directions of those deviations. Both were specified before they were run and use no arm labels.

The library-level permutation test (`perm_calibrated.py`) recomputes a statis-tic *T* for each of the 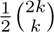 distinct splits of the 2*k* libraries into two groups of *k* (35 for *k* = 4) and reports *p* = #{*T*_split_ `≥` *T*_observed_}*/*35. The statistics are *ρ_ED_*, the pseudobulk − log_10_ of the smallest moderated-*t p* value, and the energy distance between the two library groups in the Harmony-corrected embedding. Because libraries are exchangeable under no difference and the statistics use arm labels only through the split, the test is exact. Performance is the rejection rate at the stated rule and the AUC for separating simulations with a difference from simulations with none (each approach’s own statistic). Results are in `benchmark_v2_summary.csv`, `benchmark_v2_auc.csv` and `benchmark_v2_ablation.csv`. The real-data reference ablation (`real_reference_ablation.py`) divides the all-library energy distances of GSE150949 (20 PCs) by the mean absolute split-half distance in the matrix diagonal.

### Classification and statistical analysis

To test whether two groups can be distinguished, we trained an *ℓ*_2_-regularised logistic regression [49] on the top 20 corrected PCs, evaluated by five-fold stratified cross-validation, and report accuracy and the area under the ROC curve (AUC) from out-of-fold predictions. Significance came from a label-permutation test with *p* = (*r*+1)*/*(*M* + 1) [50], *M* = 200 in the main analysis (smallest attainable *p* `≈` 0.005) and *M* = 1,000 in the library and depth controls. Batch controls used the same procedure and were (i) the two independently sequenced Day 11 libraries against each other, (ii) Day 0 versus withdrawn and Day 0 versus re-treated, and (iii) a classifier using only log total counts, mitochondrial fraction and genes detected. All pairs were depth-matched. Because every condition is one sequencing library and we have no lane or batch identifiers, cross-validation cannot keep batches apart, and condition is confounded with library. Sensitivity to the number of PCs (5 to 50), and classification after binomial count thinning to a common depth and after Scrublet doublet removal (run per library), are provided in the repository.

Direct two-group comparisons (for example a marker’s expression in withdrawn versus re-treated cells, or depth between conditions) used the Mann-Whitney *U* test [51], chosen because single-cell expression is zero-inflated and non-normal. Even after the global depth correction, the two reversibility conditions differed in raw depth (median 1,986 versus 1,311 counts, *p <* 0.0001), whereas Day 0 versus Day 11 did not (*p* = 0.11), so a targeted correction was applied. Cells were binned into ten joint depth-quantile bins and within each bin both groups were downsampled without replacement to the smaller count. This gave two groups of 862 cells (post-matching median depths 1,593 versus 1,606, Mann-Whitney *p* = 0.92), which were used for every reversibility analysis. The same procedure gave 839 cells per group for Day 0 versus withdrawn (median depths 1,608 versus 1,601, *p* = 0.87). All quantification is based on cell-level data, with *n* per condition in Table 5.

Multiple testing was handled within three families defined by method. These were the 18 CCM *p*-values for erlotinib (Additional file 1), the 10 permutation-entropy *p*-values (Additional file 1) and the 18 transfer-entropy *p*-values (Additional file 1), using the Benjamini-Hochberg procedure at *q <* 0.05 [52]. A result in which no null draw reaches the observed value has *p* = 0.005 for *M* = 200, not *p* = 0, and *q*-values were recomputed on this basis. Pooling all 46 tests in one family would push *SOX4* and *CD24* further from the threshold. After correction the erlotinib CCM panel is reduced to *EGFR* (both directions), *SOX4* (gene-to-trajectory only) and *CYP1B1* (trajectory-to-gene only). To ask whether withdrawn and re-treated cells form two modes or a shifted continuum, we ran Hartigan’s dip test [53] on PC1 of the depth-matched reversibility set, fitted one- and two-component Gaussian mixture models to PC1 and PC2 compared by BIC, and computed the agreement between the two-component clustering and the true labels. Software used was scanpy [45], scikit-learn, skccm, the diptest Python package and Scrublet. Custom implementations of the correlation dimension, permutation entropy, transfer entropy and surrogate tests are in the code repository (Availability of data and materials).

### Software requirements

The main analysis was run with Python 3.12 using numpy, scipy, scikit-learn, pandas, matplotlib, statsmodels and diptest (software versions are given with the code). The optional raw-data pipeline for GSE149383 additionally uses scanpy, anndata, seaborn, scikit-image, scrublet, skccm and joblib.

## Additional methods

Methods for convergent cross mapping, permutation entropy, transfer entropy and surrogate tests, which support only the exploratory analyses, are in Additional file 1.

## Abbreviations

AUC: area under the receiver operating characteristic curve;
BIC: Bayesian information criterion;
CCM: convergent cross mapping;
EGFR: epidermal growth factor receptor;
GEO: Gene Expression Omnibus;
IAAFT: iterative amplitude-adjusted Fourier transform;
PC: principal component;
PCA: principal component analysis;
scRNA-seq: single-cell RNA sequencing;
TKI: tyrosine kinase inhibitor;
UMAP: uniform manifold approximation and projection;
MI: unique molecular identifier.

## Supplementary information. Additional file 1 (PDF)

Exploratory analyses of the single-library erlotinib dataset (GSE149383): correlation dimension at equal sample size, cross mapping, transfer entropy, permutation entropy, marker comparison, a summary of claims and their checks, supplementary methods, and legends of Tables S1–S7 and Figs S1–S7.

## Additional file 2 (PDF)

Simulation benchmark: ground-truth toggle-switch simulations of single-library contrasts, bimodality tests, correlation dimension and binned-series analyses; robustness of the distance ratio to the structure of the library effect; and the extended benchmark of *ρ_ED_* and library-level permutation tests against alternative approaches, with full results for two, three and four libraries per arm.

## Additional file 3 (ZIP)

Numerical results as CSV files: GSE150949 noise floor, contrasts and replicate-difference dimension, dip test, correlation dimension, trajectory distances and lineage concordance, the library distance matrices, the benchmark of *ρ_ED_* and library-level permutation tests against alternative approaches (raw and summarised simulation output), the real-data reference ablation, and the simulation results, with the scripts that produce them.

## Acknowledgements

We thank William N. William Jr., M.D. (Medical Oncologist and National Leader for Thoracic Oncology, Oncoclínicas, São Paulo, Brazil) for clinical oncology perspective and for critical reading of the manuscript. He is named with his permission and did not contribute to the analyses. We used the publicly deposited datasets GSE149383, generated by Aissa and colleagues [37], and GSE150949, generated by Oren and colleagues [17].

## Declarations

### Ethics approval and consent to participate

Not applicable. No new experiments were done, and no human participants, human samples or vertebrate animals were involved. The analysed data are public datasets generated by the authors of the original studies [17, 37].

### Consent for publication

Not applicable.

### Availability of data and materials

Both datasets analysed are public in the NCBI Gene Expression Omnibus under accessions GSE149383 [37] and GSE150949 [17]. No new sequencing data were generated. The analysis code is available on GitHub (https://github.com/wasimrazaa135-ops/pc9-single-cell-attractor-inference) and on Zenodo (https://doi.org/10.5281/zenodo.24256121), respectively. Numerical results are provided as Additional file 3.

### Competing interests

The authors declare that they have no competing interests.

### Funding

The study received no specific funding.

### Authors’ contributions

**Wasim Raza**: Conceptualization, Methodology, Soft-ware, Formal analysis, Investigation, Data curation, Writing (original draft), Writing (review and editing). **Saba Mehmood**: Formal analysis, Validation, Software, Writ-ing (review and editing). **Carsten Enderlein**: Resources, Validation, Visualization, Writing (review and editing). **Fabrício Alves Barbosa Da Silva**: Conceptualiza-tion, Supervision, Project administration, Writing (review and editing). All authors read and approved the final manuscript.

### Use of generative AI

The authors declare that no generative AI tools or AI-assisted technologies were used in the conception, data analysis, code development, writing, or editing of this manuscript. All analyses and text were produced entirely by the authors.

### Authors’ information

**Wasim Raza** received his Master of Mathematics from Institut Teknologi Sepuluh Nopember (ITS) in Surabaya, Indonesia. He is now a Ph.D. student at the Federal University of Rio de Janeiro (UFRJ), Brazil, jointly affiliated with the Nanobiosystem Program and Biomedical Engineering. His work covers mathematical modelling and simulation, artificial intelligence and bioinformatics, with a focus on quantitative computational frameworks for complex biological systems.

**Saba Mehmood** holds a Ph.D. and did post-doctoral research at Universitas Airlangga (UNAIR) in Surabaya, Indonesia. Her background is in applied mathematics, functional analysis, operator theory and mathematical modelling. She is an Assistant Professor in the Department of Mathematics at Thal University Bhakkar, Pakistan, where her cross-disciplinary work links mathematical theory with the Internet of Things (IoT) and machine learning for health analytics and predictive computational modelling.

**Carsten Enderlein** earned his Ph.D. and completed postdoctoral research in Physics. He is an Associate Professor at the Federal University of Rio de Janeiro (UFRJ), based at the Duque de Caxias Campus (GERATEC), and takes part in the Nanobiosystem Program. His research is on nan-otechnology, nanoscale materials and physical biophysics, bringing physical insight to nanoscale biological interactions.

**Fabrício Alves Barbosa da Silva** received his Ph.D. and completed postdoctoral training in Scientific Computing applied to Biological Systems, with a particular focus on cancer modelling and gene regulatory networks. He is an Associate Professor at the Oswaldo Cruz Foundation (FIOCRUZ) in Rio de Janeiro, Brazil, where he leads computational systems biology research on complex biological pathways, disease dynamics and cellular simulation.

## Notes

https://github.com/wasimrazaa135-ops/pc9-single-cell-attractor-inference

https://doi.org/10.5281/zenodo.24256121

https://www.ncbi.nlm.nih.gov/geo/query/acc.cgi?acc=GSE149383

https://www.ncbi.nlm.nih.gov/geo/query/acc.cgi?acc=GSE150949

### Competing Interest Statement

The authors have declared no competing interest.

